# CDK2 kinase activity is a regulator of male germ cell fate

**DOI:** 10.1101/595223

**Authors:** Priti Singh, Ravi K. Patel, Nathan Palmer, Jennifer K. Grenier, Darius Paduch, Philipp Kaldis, Andrew Grimson, John C. Schimenti

**Affiliations:** Cornell University College of Veterinary Medicine, Dept of Biomedical Sciences, Ithaca, NY 14853.; Institute of Molecular and Cell Biology (IMCB), Agency for Science, Technology, and Research (A*STAR), Singapore 138673, and Department of Biochemistry, National University of Singapore, Singapore 117599, Republic of Singapore.; Cornell University, Weill Cornell Medicine, Dept of Urology, New York, NY.; Cornell University, Department of Molecular Biology and Genetics, Ithaca, NY.

## Abstract

The ability of men to remain fertile throughout their lives depends upon establishment of a spermatogonial stem cell (SSC) pool from gonocyte progenitors, and also maintaining the proper balance between SSC renewal and spermatogenic differentiation throughout life. Depletion of SSCs causes infertility with a Sertoli Cell Only Syndrome (SCOS) phenotype. We previously created a mouse strain in which an inhibitory phosphorylation site (Tyr15) of Cyclin-dependent kinase 2 (*Cdk2*) was altered. Juvenile males homozygous for this allele (*Cdk2^Y15S^*) initiate the first round of spermatogenesis, which originates from prospermatogonia, but meiocytes arrest due to chromosomal defects resembling those in *Cdk2^-/-^* mice. Subsequent waves of spermatogonial differentiation and meiosis were largely absent, leading to an SCOS-like phenotype. Here, we demonstrate that *Cdk2^Y15S/Y15S^* mice possess mitotically active GFRa1^+^ SSC-like cells, but they are impaired in their ability to differentiate. Marker analysis and single cell RNA-seq revealed defective differentiation of gonocytes into SSCs. Biochemical and genetic data demonstrated that *Cdk2^Y15S^* is a gain-of-function allele causing deregulated kinase activity, and its phenotypic effects could be reversed by mutating the Thr160 positive regulatory site in *cis*. These results demonstrate that precise temporal regulation of CDK2 activity in male germ cell development and in the cell cycle is critical for long-term spermatogenic homeostasis.

## INTRODUCTION

Humans and mice are capable of reproduction throughout much of their lives due to a continuously regenerating pool of spermatogonial stem cells (SSCs) in adult testes. In mice, the progenitors of SSCs, called primordial germ cells (PGCs), arise as a group of ∼45 cells in the epiblast of 6-6.5 day old embryos (Ginsburg et al., 1990). The PGCs migrate from the epiblast (extraembryonic) region of the early embryo to the location of the future gonads (the genital ridges) while proliferating during migration. Upon their arrival in the primitive gonad at around embryonic day (E) 10.5, they continue proliferating rapidly as the gonads differentiate into either primitive ovaries or testes. These germ cells, now called gonocytes (also prospermatogonia) in males, reach a population of ∼25,000 by E13.5. Around this time, female gonocytes directly enter meiosis, while the male cells largely cease proliferation for the remainder of gestation (Sasaki and Matsui, 2008; Tam and Snow, 1981).

Shortly after birth, the prospermatogonia resume proliferation to establish the permanent pool of SSCs that will seed waves of spermatogenesis throughout adult life. Additionally, a subset of these prospermatogonia initiate a distinct prepubertal round of spermatogenesis, involving several mitotic divisions before entering meiosis at approximately postnatal (P) day 10 (Bellve et al., 1977). This first round is unique because it originates from neurogenin 3 (NGN3)-expressing prospermatogonia (Yoshida et al., 2004, 2006). During adult life, new coordinated waves of spermatogenesis derive from divisions of an individual Type A_single_ (A_s_) SSC and its progeny to form Type A_paired_, (A_pr_), A_aligned_ (A_al_), and Type B spermatogonia, and ultimately preleptotene spermatocytes that enter meiosis. Through mechanisms that are not fully understood, there is an emerging picture of heterogeneity in the SSCs, in which self-renewal potential decreases as spermatogonia chain length increases [reviewed in (de Rooij, 2017)]. Waves of spermatogonial expansion are necessarily linked to replenishment of the A_s_ SSC pool from a subset of the Type A spermatogonia. This balance between SSC renewal versus differentiation must be highly regulated, otherwise there will either be an insufficient number of cells undergoing spermatogenesis, or premature exhaustion of the stem cell pool that can cause permanent infertility. Genetic or environmental events that lead to the latter, or which compromise the generation or viability of SSCs or their progenitors, can cause SCOS, a histological phenotype categorizing a subset of patients with non-obstructive azoospermia (NOA).

Normal proliferation of cells is dependent on cell cycle regulation. Key players in this process are cyclin-dependent kinases (CDKs) and their activating partner proteins, cyclins, several of which exist in mammals. CDK activity is controlled during the cell cycle in part by the association of CDKs with cyclins. In their activated state, these complexes propel cells through various stages of the cell cycle, such as entry into and through M and S phases (Satyanarayana and Kaldis, 2009). Genetic and biochemical studies have shown that the transition between active and inactive states of CDK/cyclin complexes is governed by both the interaction with CDK-inhibitory proteins (Lim and Kaldis, 2013) and also phosphorylation or dephosphorylation events at key regulatory sites on CDKs (Cuijpers and Vertegaal, 2018; Morgan, 1995).

Though the activities of cyclins and CDKs have been studied predominantly in cultured somatic cells and single-celled eukaryotes, their roles in the germline have also been investigated (Martinerie et al., 2014; Wolgemuth and Roberts, 2010). Surprisingly, despite its broad expression in many cell types, *Cdk2* is not essential for mouse viability, yet its disruption results in male and female infertility (Berthet et al., 2003; Ortega et al., 2003). Specifically, *Cdk2^-/-^* meiocytes arrest during the pachytene stage of meiotic prophase I. This arrest is triggered by defective attachment of telomeres to the nuclear envelope, resulting in their failed or incomplete synapsis of homologous chromosomes. In turn, these defects prevent homologous recombination repair of DNA damage accumulated during early meiotic prophase (Viera et al., 2009, 2015). Despite CDK2’s presence in spermatogonia (Johnston et al., 2008; Ravnik and Wolgemuth, 1999), the SSC population apparently remains functional because mutant males produce spermatocytes (albeit destined for meiotic arrest) into adulthood. These results suggest that, as in most somatic cells, CDK2 function is not essential in spermatogonia, but it may provide redundant function in those cells and non-canonical function(s) in meiocytes related to recombination in the latter (Berthet et al., 2003; Krasinska et al., 2008). Although a spermatogonia-specific deletion of *Cdk1* has yet to be described, this kinase is required for metaphase I entry at the end of the first meiotic prophase (Clement et al., 2015). CDK1 likely acts in concert with the meiosis-specific cyclin A1, which is also required at the same stage (Liu et al., 1998). In contrast, conditional ablation of CyclinB1 (*Ccnb1*), a CDK1 binding partner, was found to block proliferation of gonocytes and spermatogonia, but spermatocytes ablated for *Ccnb1* underwent meiosis normally (Tang et al., 2017).

In an effort to identify segregating infertility alleles in human populations, we used CRISPR/Cas9 editing to generate a mouse model of a missense SNP variant (rs3087335) altering the TYR15 phosphorylation site of CDK2 (Singh and Schimenti, 2015). Surprisingly, this allele (*Cdk2^Y15S^*) caused an SCOS-like phenotype. Additionally, unlike heterozygous nulls, *Cdk2^Y15S^* heterozygotes exhibited age-dependent testis histopathology and reduced sperm production, indicating that *Cdk2^Y15S^* is a gain-of-function, semidominant, allele (Singh and Schimenti, 2015). *In vitro* studies have shown that Tyr15 phosphorylation, typically catalyzed by the WEE1 kinase, negatively regulates CDK activity and thus, cell cycle progression (Gu et al., 1992; Welburn et al., 2007). We speculated that the *Cdk2^Y15S^* allele was hyperactive by virtue of being refractory to negative regulation by WEE1 (Hughes et al., 2013; Zhao et al., 2012), thus driving excessive spermatogonial proliferation and/or differentiation over SSC regeneration and maintenance, and ultimately causing SCOS.

Here, we report that the apparent SCOS phenotype in *Cdk2^Y15S/Y15S^* testes is not due to an absence of germ cells; rather, SSC-like cells are present and can divide, but their progeny fail to differentiate and subsequently are lost before entering meiosis. The germ cell defects are first detectable at postnatal day 3, where the gonocyte-to-spermatogonia transition (GST) appears disrupted as determine by single cell (sc) RNA-seq analyses. We provide evidence that CDK2^Y15S^-containing cells have altered kinase activity, and that this defect underlies phenotypes observed in such cells. Finally, we find that the disrupted regulation of CDK2 does not affect the first round of spermatogenesis, which arises from prospermatogonia. This study highlights the importance of precise regulation of CDK kinase activity in maintaining testis homeostasis.

## RESULTS

### Ablation of the TYR15 negative phosphoregulatory site in CDK2 disrupts gonocyte and spermatogonia differentiation

As summarized above, *Cdk2^Y15S/Y15S^* adult testes lacked evidence of spermatogenesis and were essentially devoid of cells positive for DDX4 (hereafter called MVH, mouse vasa homolog). MVH marks gonocytes and all juvenile germ cells, but in adults, is highly expressed in differentiated germ cells (zygotene spermatocytes and beyond) (Toyooka et al., 2000). Our working hypothesis was that since TYR15 phosphorylation by the WEE1 kinase negatively regulates CDK2 activity, then the mutant *Cdk2* allele might promote proliferation of most gonocytes in the initial spermatogenic wave, leaving the adults devoid of a renewable stem cell pool. This hypothesis predicts that mutant testes should initially have a normal gonocyte pool, but that the spermatogonial stem cells (SSCs) and differentiated progeny should become exhausted in early adulthood.

To test this hypothesis, we first quantified gonocytes in neonatal testes. The number of MVH^+^ cells in P0 *Cdk2^Y15S/Y15S^* testes was no different than in control littermates (Fig. 1A-B), indicating that the loss of germ cells occurred not during gestation, but during postnatal development. We next performed immunohistochemical analysis of mutant adult seminiferous tubule sections, which lack ongoing spermatogenesis, for the presence of rare germ cells. Remarkably, P180 *Cdk2^Y15S/Y15S^* sections contained ample numbers of cells positive for LIN28, a marker of Type A_s_ through Type B spermatogonia (Fig. 1C; Fig S1B,C), demonstrating that although mutant testes had an SCOS-like appearance, there were indeed spermatogonia present; however, they apparently were not proliferating or differentiating in a normal manner.

**Figure 1:**
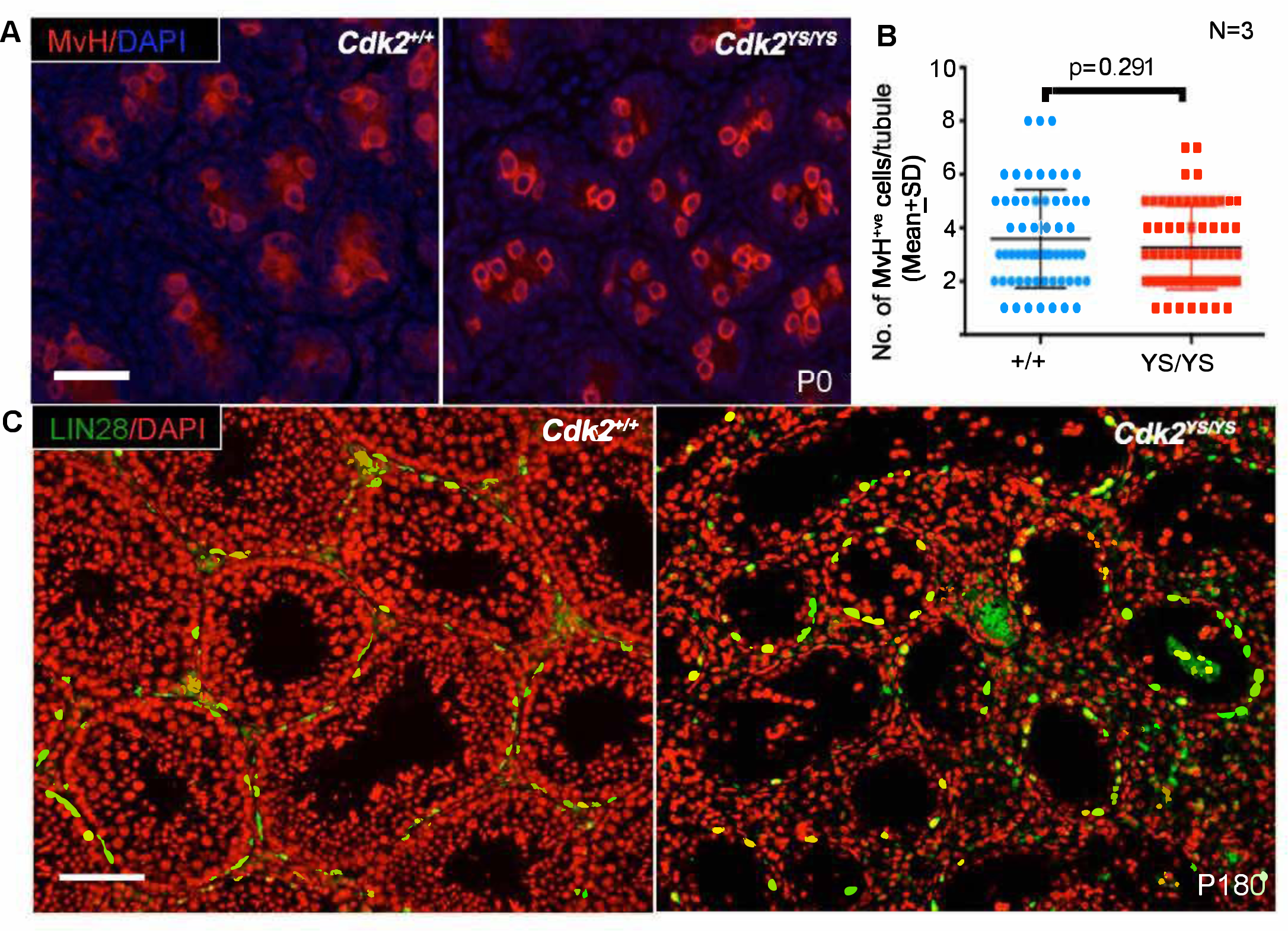
*Cdk2^YS/YS^* mice are born with a normal spermatogonial complement, but defective differentiation manifests in adulthood. **(A)** Immunohistochemical staining for MVH (red), marking germ cells in P0 sections of WT and *Cdk2^Y15S/Y15S^* testis. **(B)** Quantification of MVH^+^ germ cells per tubule cross-section in WT and *Cdk2^Y15S^* (“YS”) mutant testes (n= 3 for each group). **(C)** Immunohistochemical staining for LIN28 (green) on P180 testes for the indicated genotypes. Nuclei were counterstained with DAPI (blue in A; pseudocolored red in C). Scale bars 50 µM in (A) and 100 µm in (C).

This result led us to posit that the TYR15>SER change in CDK2 disrupts the balance between self-renewal and differentiation of SSCs in one of two ways: 1) it prevents SSC differentiation, and/or 2) it causes SSCs to permanently enter a quiescent G_0_ cell cycle stage. To distinguish between these possibilities, we characterized maturation of SSCs using the following antibodies that mark gonocytes and progressively more differentiated spermatogonia: FOXO1 (Forkhead box O1), GFRA1 (glial cell line derived neurotrophic factor family receptor alpha 1, the cell surface receptor for GDNF), PLZF (zinc finger and BTB domain containing 16, formally ZBTB16) and STRA8 (stimulated by retinoic acid 8, the expression of which coincides with meiotic entry (Kojima et al., 2019)). Whereas GFRA1 mainly labels A_s_ and A_pr_ spermatogonia, PLZF and LIN28 are also expressed in A_al_ spermatogonial chains (Tseng et al., 2015). FOXO1 transits from the cytoplasm to the nucleus as gonocytes differentiate into spermatogonia in neonatal gonads (Goertz et al., 2011). At P0, mutant gonads retained normal cytoplasmic localization of FOXO1 (Fig. S1A). We further quantified PLZF^+^, LIN28^+^ and FOXO1^+^ cells in seminiferous tubules at different times after birth (from P3-P90) by immunolabeling of testis sections. Interestingly, there were fewer spermatogonia positive for these 3 markers in mutants at P3, but not at P15 for LIN28 and only minimally for PLZF (Fig. S1B, C). In conjunction with the finding that mutants are born with normal numbers of germ cells, these results suggest either a delay or defect in differentiation of *Cdk2^Y15S/Y15S^* gonocytes to SSCs, or that those cells that progressed from gonocytes to SSCs have a lower propensity for differentiation.

To address this issue, we examined the intracellular location of FOXO1 over time. Unexpectedly, whereas all FOXO1^+^ cells in P30 WT seminiferous tubules showed nuclear localization, >40% of mutant cells exhibited cytoplasmic FOXO1 at P30; this fraction declined to ∼10% at P90 (Fig. 2A-C; Fig. S1B). These results suggest a severe defect or delay in the GST. Next, we performed a series of studies on seminiferous tubule whole mounts to determine if there were disruptions to the normal patterns of Type A spermatogonia subtypes (e.g., A_s_ vs A_pr_ vs A_al_). One model for rodent spermatogonia differentiation is that progenitor spermatogonia (SSCs) are a subset of A_s_ cells that can divide to produce and A_pr_ and A_al_ chains connected by intercellular bridges, and that cells in longer chains have decreased ability to reconstitute the SSC population (Nakagawa et al., 2010). Our studies revealed that almost all A_s_ and A_pr_ PLZF^+^ SSCs from P90 *Cdk2^Y15S/Y15S^* mice also expressed GFRA1, similar to age-matched WT controls. Mutants also contained chains of 4-8 PLZF^+^LIN28^+^ cells, however, they lacked longer chains (A_al16-32_) that are typical in WT testes (Fig. S2A) (Buaas et al., 2004; Zheng et al., 2009). This result is consistent with the reduced PLZF^+^ and LIN28^+^ spermatogonia in testes cross sections of young (P5) and old (P90) *Cdk2^Y15S^* homozygotes (Fig. S1B, C).

**Figure 2:**
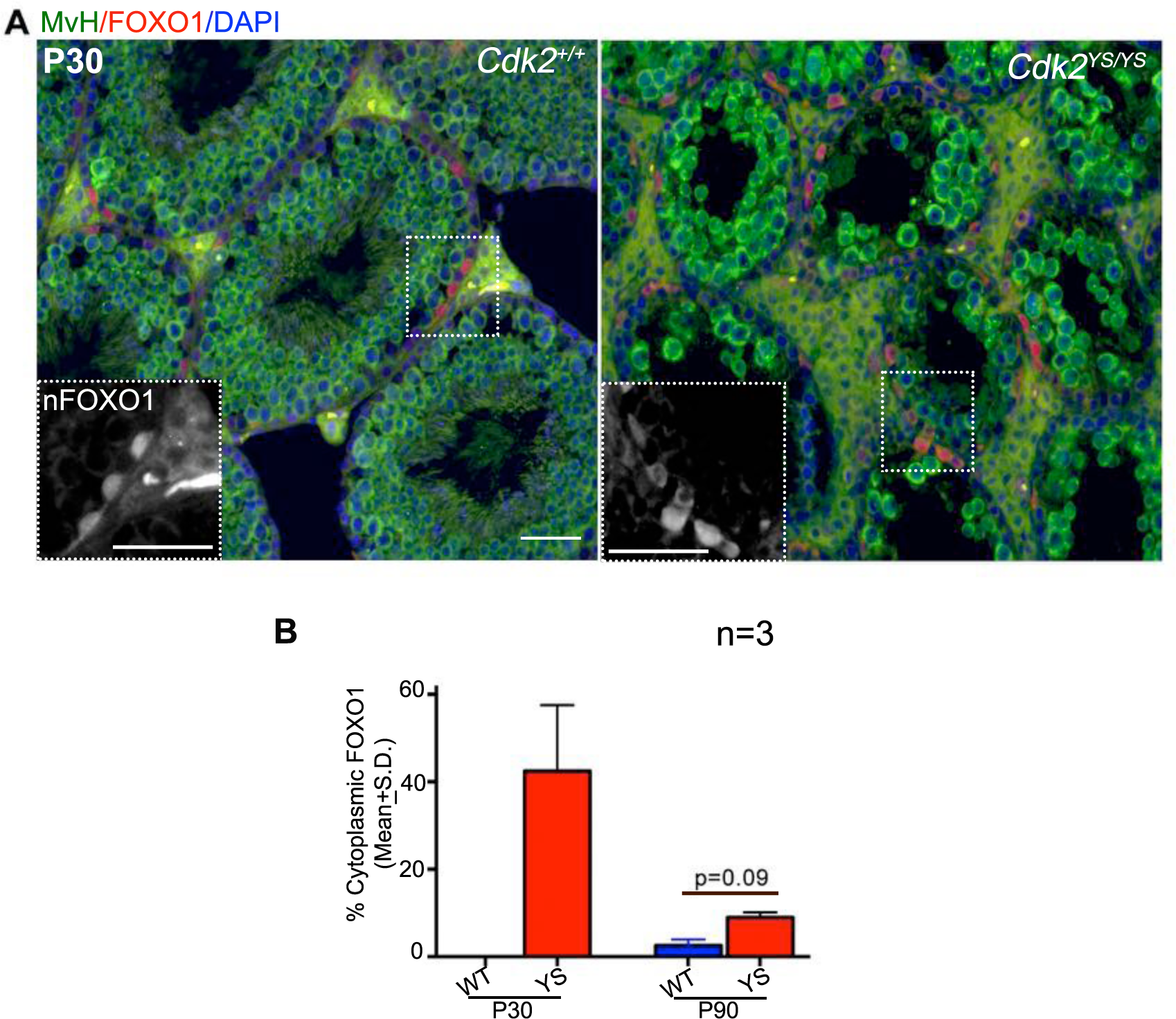
*Cdk2^YS/YS^* gonocytes do not differentiate normally into SSCs. **(A)** Immunolabeling of P30 testes histological sections labeled by MVH (green) and FOXO1 (red) showing presence of cFOXO1^+^ (cytoplasmic FOXO1) germ cells which localized in the nucleus (nFOXO1) in WT tubules. **(B)** Percentage of cFOXO1 in P30 and P90 seminiferous tubule cross sections showing massive increase in cFOXO1^+^ cells in *Cdk2^Y15S^* (YS) homozygotes. Scale bar, 50 μm.

To explore the basis for the defects in spermatogonial distributions, we compared the distribution and frequency of GFRA1^+^ progenitor spermatogonia in adolescent (P15, during the first round of spermatogenesis) vs adult (P90) tubules that lack spermatogonial differentiation. *Cdk2^Y15S/Y15S^* mutants had about twice as many GFRA1^+^ cells at both ages compared to WT (Fig. 3A-B). More striking was that mutant tubules contained long chains of GFRA1^+^ spermatogonia (referred to as ‘GFRA1^+^ A_al>6_’) in P15 tubules, which were absent in WT (Fig. 3A-D). Hypothesizing that these GFRA1^+^ A_al>6_ cells might be in an abnormal, delayed state of differentiation, we examined these chains for co-expression of SALL4, a marker of differentiation-primed spermatogonia (Nakagawa et al., 2010). Interestingly, the abnormal GFRA1^+^ A_al_ chains in mutants expressed SALL4 (Fig. S2B). The GFRA1^+^ A_al_ chains disappeared by P90, yet mutant tubules had more GFRA1^+^ A_s_ and A_pr_ cells compared to WT at this age (Fig. 3B-C). The combined data indicate that the *Cdk2^Y15S^* allele causes a defect in differentiation of gonocytes and GFRA1^+^ progenitor spermatogonia.

**Figure 3:**
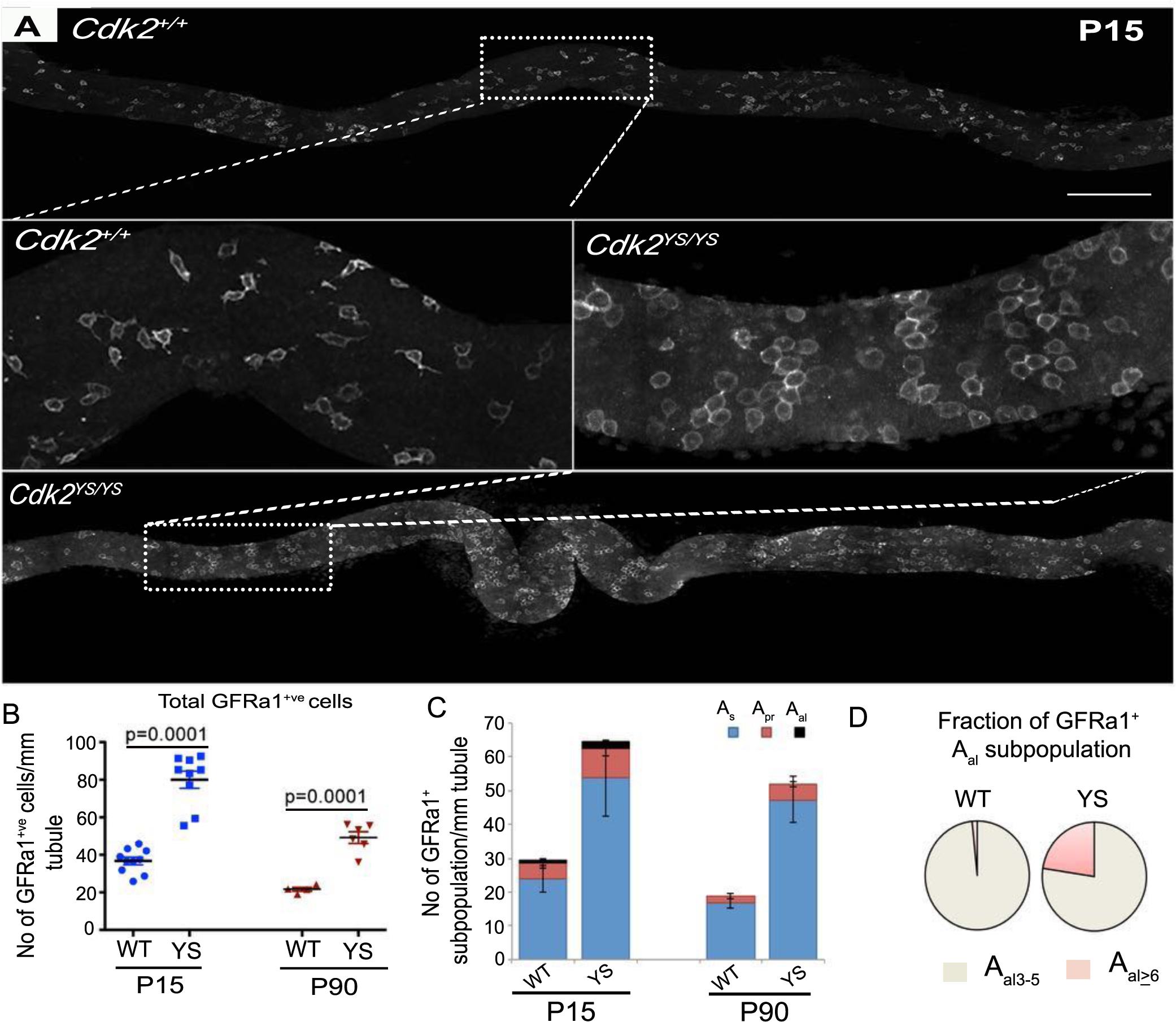
*Cdk2^YS/YS^* seminiferous tubules contain more undifferentiated spermatogonia. **(A)** Representative confocal (stitched) images of P15 seminiferous tubules of the indicated genotypes, showing density of GFRa1^+^ spermatogonia (white). Images were captured along the length of seminiferous tubules and the outermost 2-3 optical sections are projected. Insets show higher magnification of the boxed areas. Scale bar 200 µm. Density of **(B)** total and **(C)** A_s_, A_pr_ and A_al_ GFRa1^+^ spermatogonia in seminiferous tubules of P15 and P90 testes of indicated genotypes. n ≥ 3 in either genotype (except n=2 for P90 WT). 2 or more tubules (>8 mm long) were analyzed in each animal. **(D)** Pie chart showing distribution of GFRa1+ A_al3-5_ and A_>6_ subpopulation in P15 *Cdk2^+/+^* and *Cdk2^YS/YS^* gonads.

### Regulation of CDK2 activity is critical for balancing spermatogonia progenitor self-renewal vs differentiation

Whereas SSC differentiation or maintenance was not obviously impacted in *Cdk2* null mice, *Cdk2^Y15S^* heterozygotes exhibited an age-related decrease in spermatogenesis. This suggests that *Cdk2^Y15S^* is a hypermorphic, or a gain-of-function allele. A classical model of SSC homeostasis posits that an A_s_ spermatogonium either self-renews by forming two new A_s_ spermatogonia or takes a step towards differentiation by forming an A_pr_ two-cell chain (E. F. Oakberg, 1971). Given the increase in GFRa1^+^ spermatogonia in the *Cdk2^YS/YS^* testes (Fig. 3), we hypothesized that *Cdk2^Y15S^* either drives abnormal proliferation, and/or it skews these cells towards self-renewal instead of differentiation.

To test this, we assayed proliferation of spermatogonia by pulse labelling with the DNA analog EdU. P2 males were injected with EdU, sacrificed 4 hours later, then the testes were immunolabeled for PLZF^+^. In both mutants and WT, >99% of PLZF^+^ cells were negative for EdU (not shown), consistent with this being the period before gonocytes exit mitotic arrest to begin establishing the spermatogonial pool (Yang and Oatley, 2014). However, in older animals, we noticed severe proliferation defects in GFRa1+ and FOXO1+ cells, representing undifferentiated states. At P15, there were ∼1.6 fold more proliferating (EdU+) GFRa1^+^ cells in mutant homozygotes than WT, and this disparity persisted through P90 (Fig. 4C). Furthermore, the fractions of replicating A_s_ and A_pr_ GFRa1^+^ spermatogonia were higher in mutant than WT, and most dramatically, the A_al>6_ GFRa1^+^ category was unique to the mutant (Fig. 4B, D). Similarly, P90 *Cdk2^Y15S/Y15S^* testes had more proliferating FOXO1+ (19%) cells (Fig. 4F). In contrast, P90 testes had ∼40% fewer proliferating LIN28^+^ germ cells, which include more differentiated A_s_ spermatogonia (Fig. 4E). Collectively, our results support the notion that normal homeostasis of the stem cell niche is disrupted in *Cdk2^Y15S/Y15S^* mice, causing elevated proliferation of SSCs without normal differentiation.

**Figure 4:**
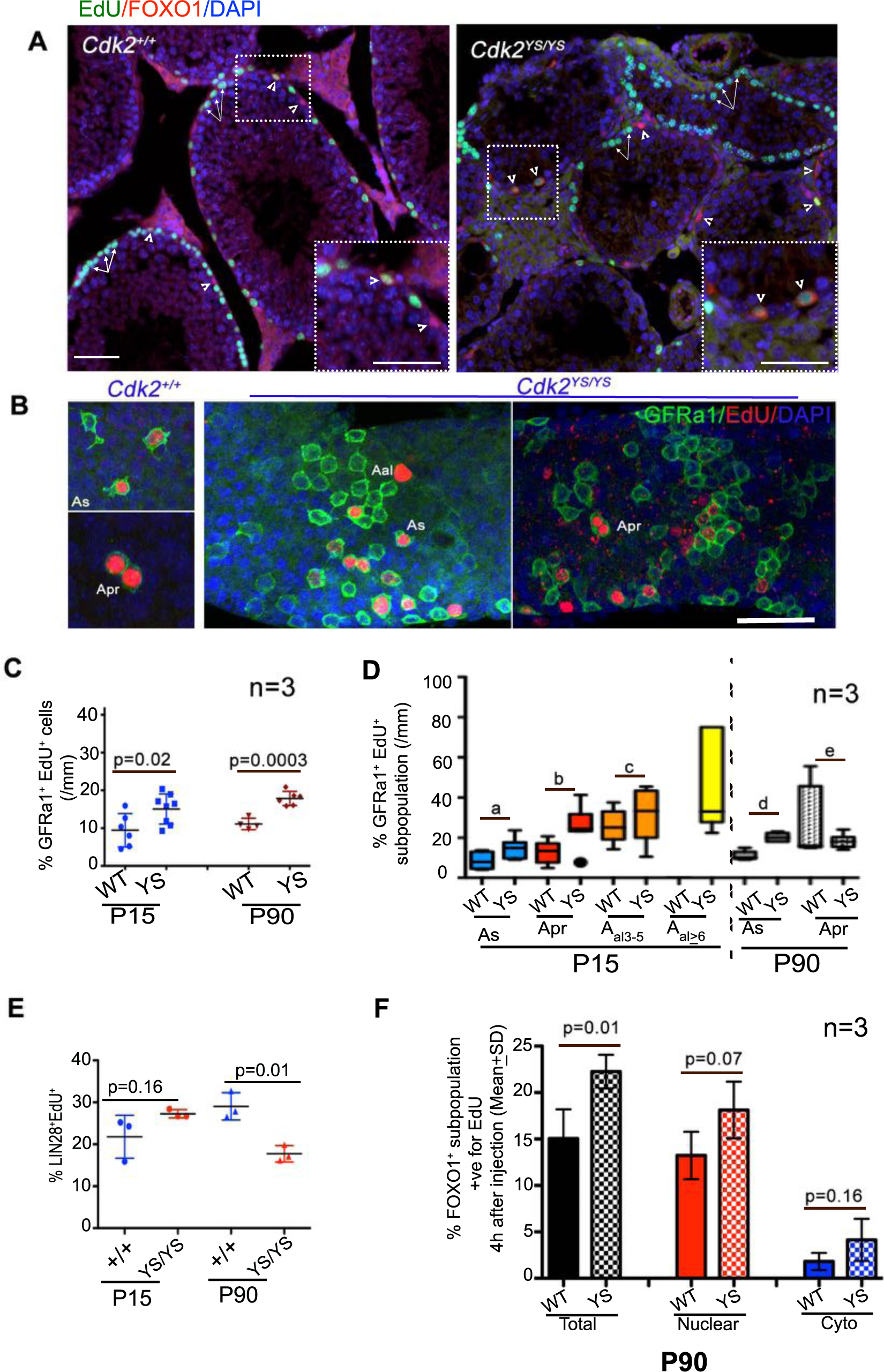
Mutation of the CDK2^Tyr15^ phosphorylation site impedes differentiation of GFRa1^+^ A_s_ spermatogonia. **(A)** Assessment of DNA replication in FOXO1^+^ germ cells. P30 mice were pulsed with EdU for 4h, then labeled for EdU incorporation and FOXO1 (red). **(B)** Representative images of whole mount seminiferous tubules (P15) showing abnormally long chains GFRa1^+^ and EdU^+^ spermatogonia. Mice were injected with EdU for 4 hours prior to sacrifice and processing of testes. The data are quantified in **(C). (D)** Quantification of A_s_, A_pr_ and A_al_ spermatogonia that retain EdU label at time of testis recovery (mean ± SD, n ≥ 3 for each group). p-values are as follows: a=0.03, b=0.01, c=0.45, d=0.0001, e=0.38. **(E)** Quantification showing percentages of replicating LIN28^+^ spermatogonia**. (F)** Percentage of replicating FOXO1^+^ cells at P90. Greater than 100 and 55 tubules were analyzed from three different WT and mutant animals, respectively. Scale Bar = 50 μm in A and B.

Despite the evidence for spermatogonial cycling in the absence of differentiation, the seminiferous tubules never (up to 20 months of age) became replete with undifferentiated cells, a condition that might lead to, or resemble, tumorigenesis. Therefore, we hypothesized that such hyperproliferating progenitor spermatogonia were being eliminated by apoptosis. Indeed, there were nearly 6-fold more TUNEL^+^ seminiferous tubules containing 2-4 fold more apoptotic germ cells in mutants vs WT at P30 (Fig. S3A-C). Furthermore, double immunolabeling for FOXO1 and cleaved PARP revealed the presence of spermatogonia undergoing apoptosis in mutants (Fig. S3D). These combined results suggest that *Cdk2^Y15S/Y15S^* undifferentiated progenitor GFRa1^+^ spermatogonia enter S phase normally but were incapable of differentiating; instead, they appear to proliferate, accumulate, and eventually undergo apoptosis.

### Single cell transcriptome analysis reveals defects in differentiation of Cdk2^Y15S/Y15S^ gonocytes and SSCs

The germ cell population at birth is characterized by substantial functional and molecular heterogeneity (Culty, 2013). Gonocytes in P3 testes, which constitute about half of all germ cells (Ohmura et al., 2004), can undergo one of three immediate fates: 1) re-enter the cell cycle, 2) migrate towards the basement membrane of the seminiferous cords to become SSCs, or 3) differentiate into spermatogonia that seed the first wave of spermatogenesis. Based on OCT4 expression, gonocytes gradually disappear during the first week of life, decreasing from 98% to 30% of all germ cells (Ohmura et al., 2004). Thus, the first postnatal week lies within the GST interval [although the timing may differ in Swiss outbred mice (Pui and Saga, 2017)]. Based on our findings that *Cdk2^Y15S^* mutants are born with a normal number of germ cells, and the first apparent abnormality was a deficit of PLZF^+^/ LIN28^+^/ FOXO1^+^ spermatogonia at P3, we hypothesized that the neonatal germ cell pool was defective in differentiating into SSCs.

To gain insight into the molecular and cellular defects in *Cdk2^Y15S^* mutants, we performed scRNA-seq on unsorted cells from WT and mutant P3 testes using the 10X Genomics platform. Data were obtained from WT (5,061 cells), *Cdk2^Y15S/+^* (4,958 cells), and *Cdk2^Y15S/Y15S^* (4,403 cells) testes. There was a median of 2,239 genes and 5,751 mRNA molecules detected per cell. Somatic and germ cells were evident in the clustering analysis of 14,422 cells from testes across all genotypes. Following unbiased k-means clustering of cells based on gene-expression differences, we identified 5 major cell clusters (Fig. 5A). The cells types within each cluster were identified by markers diagnostic (Fig. S4) of particular gonadal lineages. The percentages and numbers of cells in each of the 5 clusters were as follows: MVH^+^ germ cells (2.06%, n=299); WT1^+^ Sertoli cells (46.88%, n=6,781); CYP11a1^+^ Leydig cells (1.5%, n=217); MYH11^+^ myoid cells (22.15%, n=3,204); and VCAM1^+^ peritubular/epithelial cells (27.9%, n=3,963).

**Figure 5:**
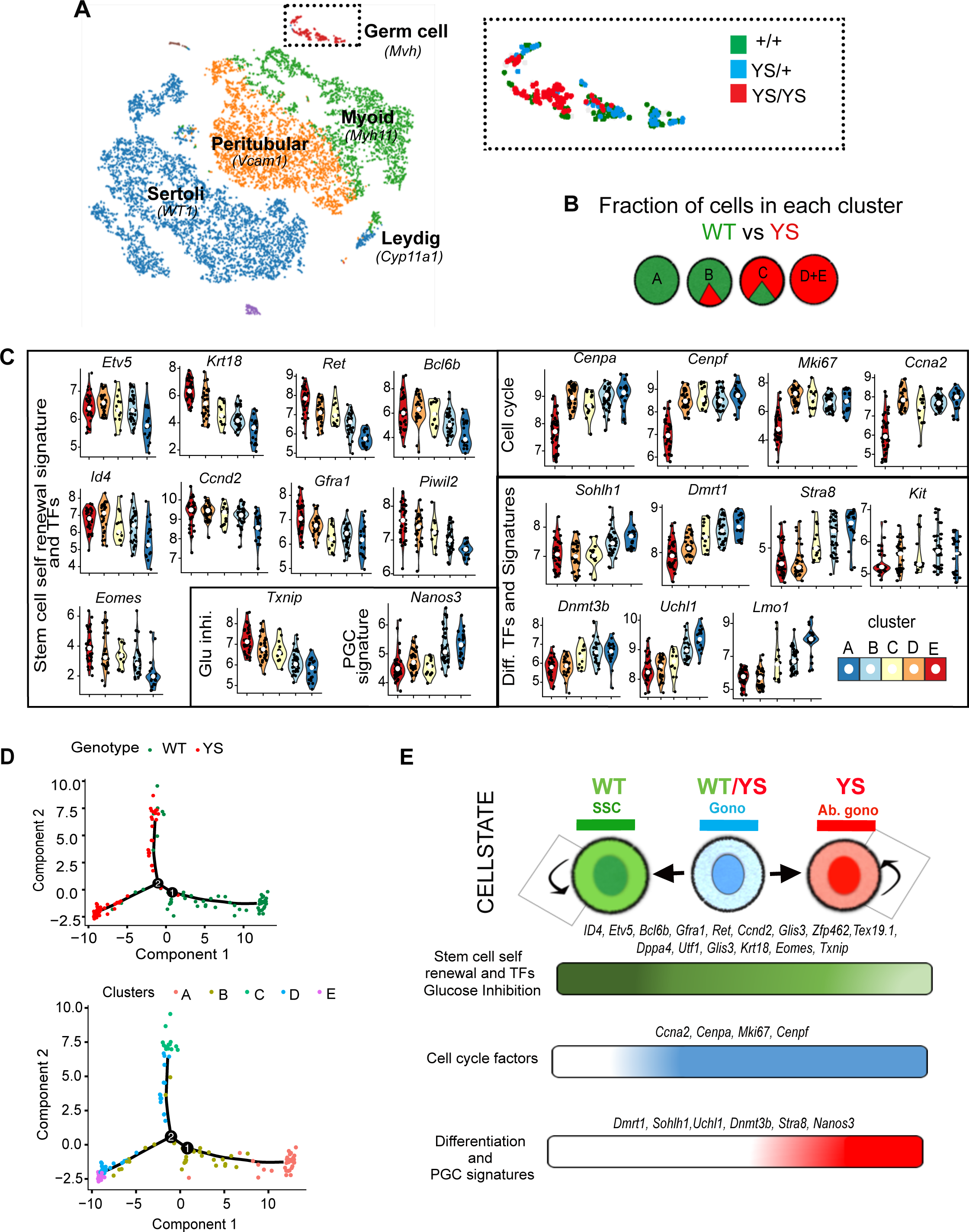
Single cell RNA sequencing of P3.5 testes reveals abnormal differentiation of *Cdk2^Y15S/Y15S^* gonocytes. **(A)** K-means clustering analysis of 14,422 testicular cells based on single-cell transcriptome data. Cell types were classified based on expression of indicated key marker genes. Inset shows the MVH^+^ germ cell cluster consisting of 263 total cells (WT = 106, *Cdk2^Y15S/Y15S^* = 58, *Cdk2^Y15S/+^* = 99), color coded by genotype. **(B)** Clustering of cells by genotype. **(C)** Violin plots showing log_2_-transformed expression levels (reads per million (RPM)) of key marker genes in each cluster. **(D)** Monocle based pseudotime trajectory analysis. Plots are based on SAVER-imputed datasets. **(E)** Schematic summary of cell states and transitions. Key gene expression clusters define three distinct cellular states in WT and *Cdk2^Y15S/Y15S^* cells. SSC= Spermatogonial stem cells; Gono.= Gonocytes; Ab. gono. = Abnormal gonocytes. YS, *Cdk2^Y15S/Y15S^*.

To characterize potential GST defects in the mutant, we focused on the MVH^+^ germ cell population. Interestingly, this group of cells consisted apparently of two populations that differed dramatically (> 2-fold) in the number of unique transcripts (UMIs) per cell (Fig. S5A). This broad bimodal distribution was not observed in somatic cell populations. We considered two potential explanations for this observation. One is that the population with higher UMIs represents cell doublets, and is thus an artifact. The confounding effects of doublets in SC data is often resolved by excluding those cells bearing higher UMIs (Ziegenhain et al., 2017). A second possibility is that neonatal germ cells are transcriptionally more active than somatic cells, which is an inherent property of embryonic germ and stem cells (Percharde et al., 2017). To test the first possibility, we used two algorithms for predicting doublets from single-cell expression data: DoubletFinder (data not shown) (McGinnis et al., 2018) and DoubletDetection (https://github.com/JonathanShor/DoubletDetection). Outputs from both algorithms indicated that the germ cells with higher UMIs (>10,000) are no more likely to be doublets than those with lower UMIs (Fig. S5B). To address the second possible explanation, we quantified total RNA from equal numbers of FACS-isolated *Oct4*-GFP+ cells from neonatal testes of transgenic mice (Szabó et al., 2002). OCT4 is a pluripotency marker and its expression is restricted to undifferentiated, prepubertal prospermatogonia and gonocytes (Ohbo et al., 2003; Szabó et al., 2002). This revealed 7-fold, 8.3-fold, and 2.4-fold higher amounts of RNA in GFP^+^ germ cells compared to lung, brain, and GFP-testis (presumably somatic) cells, respectively (Fig. S5D). To identify the source of cell barcodes with lower UMI counts, we performed a knee-plot analysis, which is used to determine the quality of a cell’s gene expression profile (Macosko et al., 2015). The analysis revealed that the cells with low UMI counts fall below the inflection point (“knee”) in the curve, suggesting that these cells are likely technical artifacts, as commonly observed in scRNA-seq experiments (Fig S5C). Taken together, these data indicate that the majority of germ cells have more transcripts (>10,000) than somatic cells (right half of the bimodal distribution, Fig. S5A), which represented 65%, 69% and 73% of the MVH^+^ germ cell populations in WT, *Cdk2^Y15S/+^* and *Cdk2^Y15S/Y15S^* samples, respectively. Moreover, a subset of the germ cells (left half of distribution) likely derive from poor-quality profiling. Since *Cdk2^Y15S/+^* clusters closely overlapped with those from WT, subsequent analyses utilized only cells from WT and *Cdk2^Y15S/Y15S^* that had elevated UMI counts (>10,000) (Islam et al., 2014). After quality control and data filtering using Seurat (Butler et al., 2018), we identified 10,451 confidently quantified genes from both genotypes (see Methods).

Since only a fraction of all transcripts are detected in scRNA-seq, the resulting expression matrices are sparse. To compute estimates for the missing data, we employed SAVER (single-cell analysis via expression recovery) (Huang et al., 2018), which uses information across all genes and cells in a dataset to impute gene expression values. We identified five subgroups of MVH^+^ germ cells, designated A-E (Fig. 5B), using hierarchical clustering based on the expression of the most divergently expressed genes (n=734; see methods) (Fig. S6). As depicted in Fig. 5B, cluster A was exclusive to the WT sample, whereas D and E were exclusive to the *Cdk2^Y15S/Y15S^* sample. Clusters B and C were preferentially enriched in WT and the *Cdk2^Y15S/Y15S^* samples, respectively.

Clusters A and B express several well-characterized markers of undifferentiated germ cells, such as *Gfra1* and *Id4* (Fig. 5C). Based on the following distinctions between the two groups, we tentatively classified cluster A as SSCs (A_SSC_) and cluster B as gonocytes (B_Gono_). Cluster A was relatively depleted in cell cycle factors such as *Ccnb1* and *Cenpf* (Fig. 5C), consistent with the idea that ‘true’ SSCs would have a lower proliferative index. Furthermore, this cluster was enriched for the inhibitor of glucose transport *Txnip*, indicating decreased metabolism, consistent with lower proliferation. Finally, to validate the cluster identities, we combined previously published scRNA-seq datasets from flow sorted OCT4^+^ (Liao et al., 2017) and ID4^+^ cells (Song et al., 2016), and identified 50 and 99 genes uniquely expressed in gonocytes and SSCs, respectively, which we then used as diagnostic markers for cell identities (Fig. S7A,B; Table S1). GSEA analyses revealed that highly expressed genes in the SSC genesets were up-regulated in cluster A compared to the reference cluster B, whereas highly expressed genes in gonocytes showed up-regulation in cluster B (Fig. S7B). The combined data led us to conclude that cluster A consists of SSCs, and cluster B consists of gonocytes.

To better define the cellular defects in mutant germ cells present at P3, we more closely examined the expression patterns of clusters A and D/E, which are the populations most unique to WT and *Cdk2^Y15S/Y15S^,* respectively. Compared to cluster A, the following features were characteristic of D/E: 1) cell cycle signature genes and E2F targets were enriched (Figs. 5C,E; 6F); 2) SSC genes were expressed at relatively low levels (Fig. 5C); and 3) the PGC/gonocyte marker NANOS3 and differentiating spermatogonia signatures (*Stra8, Lmo1, Uchl1, Dmrt1, Sohlh1, Dnmt3b*) were coexpressed and enriched (Fig. 5C, E). Altogether, these results define unique germ cells in mutants that express an unusual combination of cell cycle, differentiation and gonocyte signatures. Importantly, when we repeated these analyses using gene expression data without SAVER-based imputation, our results were consistent (Fig. S8), suggesting that SAVER imputation of scRNA data is not compromising our findings.

To further unravel the relationship between gene expression patterns of the 5 germ cell clusters and developmental state, we performed pseudo-time analysis (Trapnell et al., 2014) on the MVH^+^ germ cells. In WT, this analysis supported a trajectory path bifurcating from cluster B (WT_gono_) to clusters A (WT_SSCs_) and C (differentiation-primed gonocytes, or WT_diff-Gono_) (Fig. 5E), with the latter being defined by virtue of retaining both gonocyte and differentiation signatures (Fig. 5C-E). This trajectory path in WT is consistent with the known developmental progression. Interestingly, only ∼8% of mutant germ cells fell into Cluster B, and these eventually differentiate into two directions: 1) a small subset towards cluster C (YS_diff. gono; ∼_12%) and 2) ∼80% towards mutant-specific clusters D+E (Fig. 5D,E). It is worth noting that trajectory analyses using non-imputed data gave a different topology. Cluster C, instead of branching out as a separate cluster, appeared as a transient stage on developmental trajectory differentiating from cluster B to cluster E (Fig. S7C,D). Nevertheless, the analyses indicate a profound defect in differentiation of mutant germ cells, and are consistent with the idea that mutant gonocytes do not undergo a normal GST.

### CDK2^Y15S^ has altered kinase activity that impacts gonocyte fate

We previously hypothesized that *Cdk2^Y15S^* is a hypermorphic allele by virtue of lacking the target (TYR15) of inhibitory phosphorylation (Singh and Schimenti, 2015). However, we considered the possibility that the mutant version (SER15) could still be phosphorylated by an unknown kinase to enable some degree of negative regulation. To test this, we expressed MYC-tagged WT (CDK2-TYR15), mutant (SER15) and also PHE15 cDNAs in HEK293T cells, performed mass spectrometry (LC-MS/MS) analysis on immunoprecipitates (Fig. S9A), then analyzed the mass:charge (m/z) spectra for evidence of phosphorylated TYR/SER/PHE15 residues in the corresponding samples. The presence of WT and mutant peptides (IGEGT**S**GVVYK and IGEGT**F**GVVYK for TYR15 and PHE15, respectively) were manually confirmed in both samples. Phosphorylation was detected only at the WT CDK2^Y15^ residue, indicating that CDK2^S15^ is not a phosphorylatable substrate, at least in cultured cells (Fig. S9B).

Next, we assayed the ability of CDK2 isolated from WT and mutant P10 spleens to phosphorylate a histone H1 substrate (testis was not used as a source due to cellularity differences between mutant and WT). Consistent with ablation of the TYR15 inhibitory phosphorylation site and previous reports examining *Cdk2^T14AY15F^* activity in mouse tissues and MEFs (Zhao et al., 2012), CDK2 immunoprecipitated from *Cdk2^Y15S/+^* spleens displayed 1.5-fold more kinase activity than WT (Fig. S10B-D). Counterintuitively, material immunoprecipitated from *Cdk2^Y15S/Y15S^* spleens had >5-fold reduced kinase activity compared to WT. This may reflect the consequence of excessive CDK kinase activity, which can be toxic to cell cycle progression in a mechanism involving p21-mediated inhibition of CDK2-Cyclin (Fig. S11; see discussion) (Hughes et al., 2013; Szmyd et al., 2019; Zhao et al., 2012).

As an orthogonal assessment of CDK2 activity in germ cells, we compared expression levels of 97 key CDK2 activity signature genes in single cell clusters defined above (Table S2) (McCurdy et al., 2017). WT_SSC_ cells (cluster A) had much lower expression of CDK2 activity signature genes compared to all other clusters including WT cells in cluster B and all *Cdk2^Y15S^* clusters (Fig. 6A). Furthermore, CDK2 kinase activity, as inferred by the median of normalized expression of CDK2 activity signature genes per cell across clusters A, B, D and E, was lowest in WT_SSCs_ and highest in mutant-specific clusters D and E (Fig. 6B,C). GSEA analysis indicated up-regulation of CDK2 activity signature genes in clusters B and E compared to cluster A, suggesting that first, cluster E is more like cluster B with respect to CDK2 kinase activity, and second, that cells in cluster E have a higher propensity to cycle than those in cluster A (Fig. 6C). Interestingly, in support of this observation, genes up-regulated in cluster E compared to cluster A (q ≤0.05; FC≥0.2) also exhibited the enrichment of cell cycle related gene ontology (GO) terms (Fig. 6D). Overall, these data indicate that removing a layer of negative regulation to CDK2 activity disrupts the normal differentiation of gonocytes and SSCs into downstream cell types.

**Figure 6:**
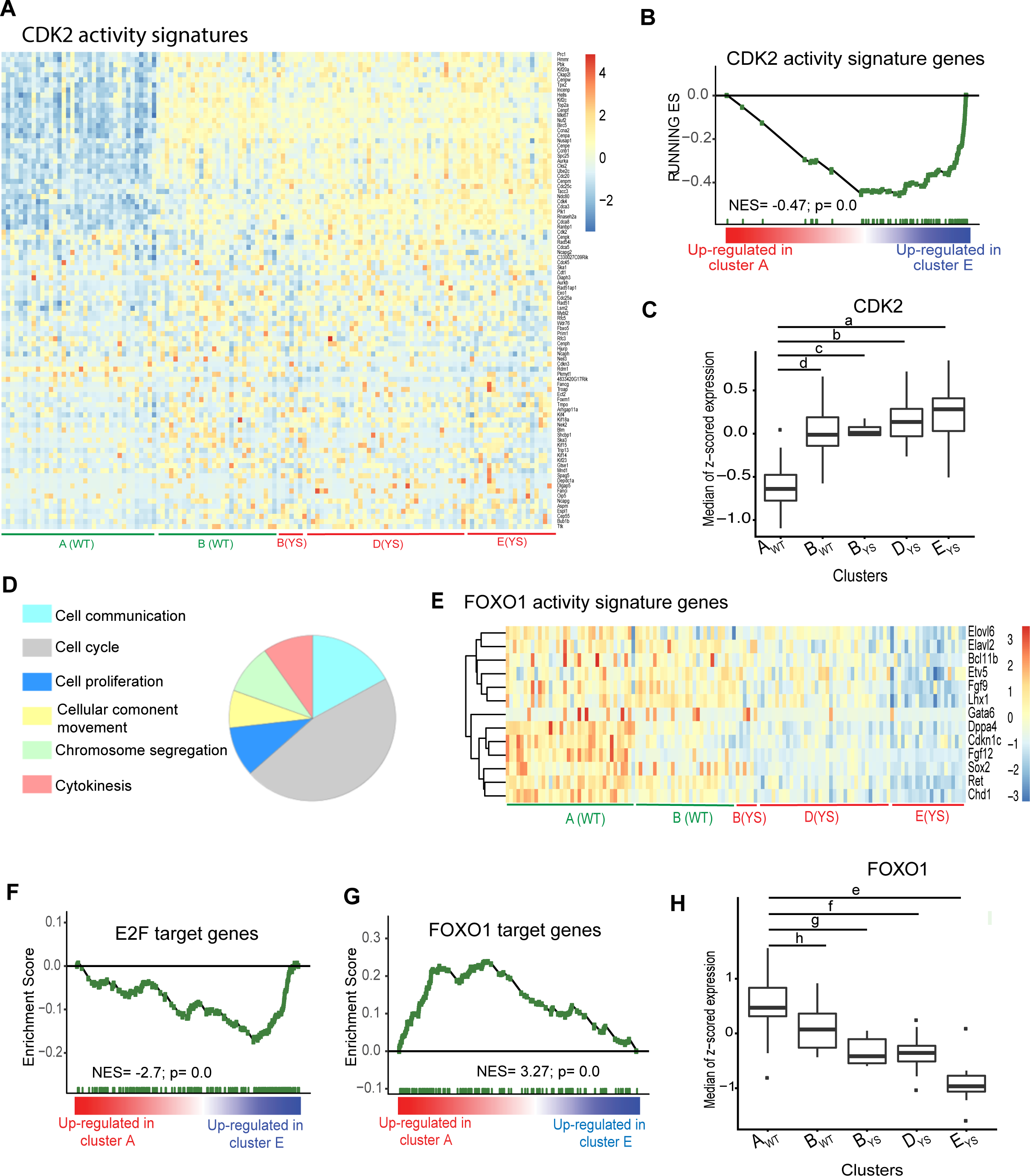
Downstream effects of impaired CDK2 activity in *Cdk2^YS/YS^* germ cells. **(A)** Heatmap showing expression of CDK2 activity signature genes (McCurdy et al., 2017) in indicated clusters of *Cdk2^+/+^* and *Cdk2^YS/YS^* germ cells. The color key is same as Fig 5A. **(B)** GSEA enrichment score plots for CDK2 activity signature genes in clusters A vs E comparisons. **(C)** CDK2 activity scores defined as median of normalized expression of CDK2 target genes per cell, across indicated clusters. (Student’s t-test p-values: a= 5.0×10^-11^; b= 4.5×10^-12^; c= 5.7×10-19; d= 1.2×10^-11^). YS = *Cdk2^Y15S/Y15S^*. **(D)** Enrichment analysis for GO-slim “biological process” terms in genes up-regulated in cluster A compared to cluster E. **(E, F)** GSEA enrichment score plots for E2F and FOXO1 target-genes (MSigDb) in cluster A vs E comparison. **(G)** Heatmap showing expression of FOXO1 targets in indicated germ cell clusters and genotypes. **(H)** FOXO1 activity defined as median of normalized expression of FOXO1 target genes per cell, across indicated clusters. (Student’s t-test p-values: e= 8.9×10^-05^; f= 6.2×10^-05^; g= 1.5×10^-13^; h= 3.7×10^-19^).

The potential molecular basis for this developmental disruption likely stems from alteration of two known functions of active CDK2: 1) phosphorylating the RB1 transcriptional repressor to inactivate it, enabling timely induction of the E2F transcription factors (TFs) to drive transition to S phase (Morris et al., 2000); and 2) inhibiting cytoplasmic-to-nuclear localization of the FOXO1 TF (nFOXO1 is essential for SSC maintenance) (Huang et al., 2006)(Goertz et al., 2011). If CDK2 is indeed controlling the gene regulatory network via E2F and FOXO1, then levels or activity of their downstream targets would be expected to be affected in *Cdk2^Y15S/Y15S^* cells. Consistent with this conjecture, GSEA analysis revealed that FOXO1 targets are up-regulated and highest in cluster A compared to cluster E, where expression is lowest (84 genes; nES: 3.27; FDR: 0) (Fig. 6 G,H). In contrast, E2F target genes are up-regulated in cluster E (Fig. 6F; 66 genes; nES: −2.7; FDR: 0). These results indicate that FOXO1 activity is significantly greater in cluster A than E (Student’s t test, p = 3.7×10^-19^) (Fig. 6H).

### Phosphorylation states at Tyr15 and Thr160 residues control CDK2 activity in male germ cells

If disrupting the ability to negatively regulate CDK2 in adult GFRa1^+^ SSCs favors cell cycle progression over differentiation, then a compensatory alteration that dampens or eliminates CDK2 activity might counteract the aberrant phenotype of *Cdk2^Y15S^* mutant spermatogonia. To test this, we used CRISPR/Cas9 to mutate threonine 160 to alanine (T160A) in the *Cdk2^Y15S^* allele. Phosphorylation of Thr160 is required for activation of CDK2 (Gu et al., 1992; Kaldis, 1999). Biochemical analyses of immunoprecipitated CDK2^T160A^ revealed that this is a “kinase dead” allele that causes infertility in both sexes, albeit with less severe meiotic phenotypes than in *Cdk2^-/-^* mice (Chauhan et al., 2016). Mice bearing the doubly mutated *Cdk2* allele (*Cdk2^Y15S,T160A^*, abbreviated as *Cdk2^YS-TA^*) were then produced, bred to homozygosity, and phenotypically analyzed with respect to spermatogenic progression in adults.

We observed three striking phenotypic differences between the *Cdk2^Y15S^* and *Cdk2^YS-TA^* alleles in adult (P120) males. First, whereas *Cdk2^Y15S^* heterozygotes had small testes and a markedly reduced sperm count as previously reported (Singh and Schimenti, 2015), *Cdk2^YS-TA^* heterozygotes were indistinguishable from WT and *Cdk2^+/-^* controls in both respects, and thus resemble *Cdk2^+/-^* mice (Fig. 7A-C). Second, as with null mice but unlike *Cdk2^Y15S^*, *Cdk2^YS-TA^* homozygotes females were sterile (n=3 in testing with WT males). Third, although *Cdk2^YS-TA/YS-TA^* adult males were severely hypogonadal (Fig. 7A-B) and azoospermic, similar to *Cdk2^Y15S/Y15S^*, they exhibited differentiating spermatogonia and meiocytes, which were completely missing from age-matched *Cdk2^Y15S^* homozygotes (Fig. 7D). At this level of analysis, the *Cdk2^YS-TA^* allele resembles the null phenotype, which is characterized by meiotic prophase I arrest in males and infertility in both sexes (Viera et al., 2009) (Chauhan et al., 2016). Collectively, our results imply that the failed spermatogonial differentiation phenotype caused by the *Cdk2^Y15S^* allele is a result of altered kinase activity from abolition of a WEE1 phosphorylation site. As a consequence of this defective negative regulation, this allele acts semidominantly.

**Figure 7:**
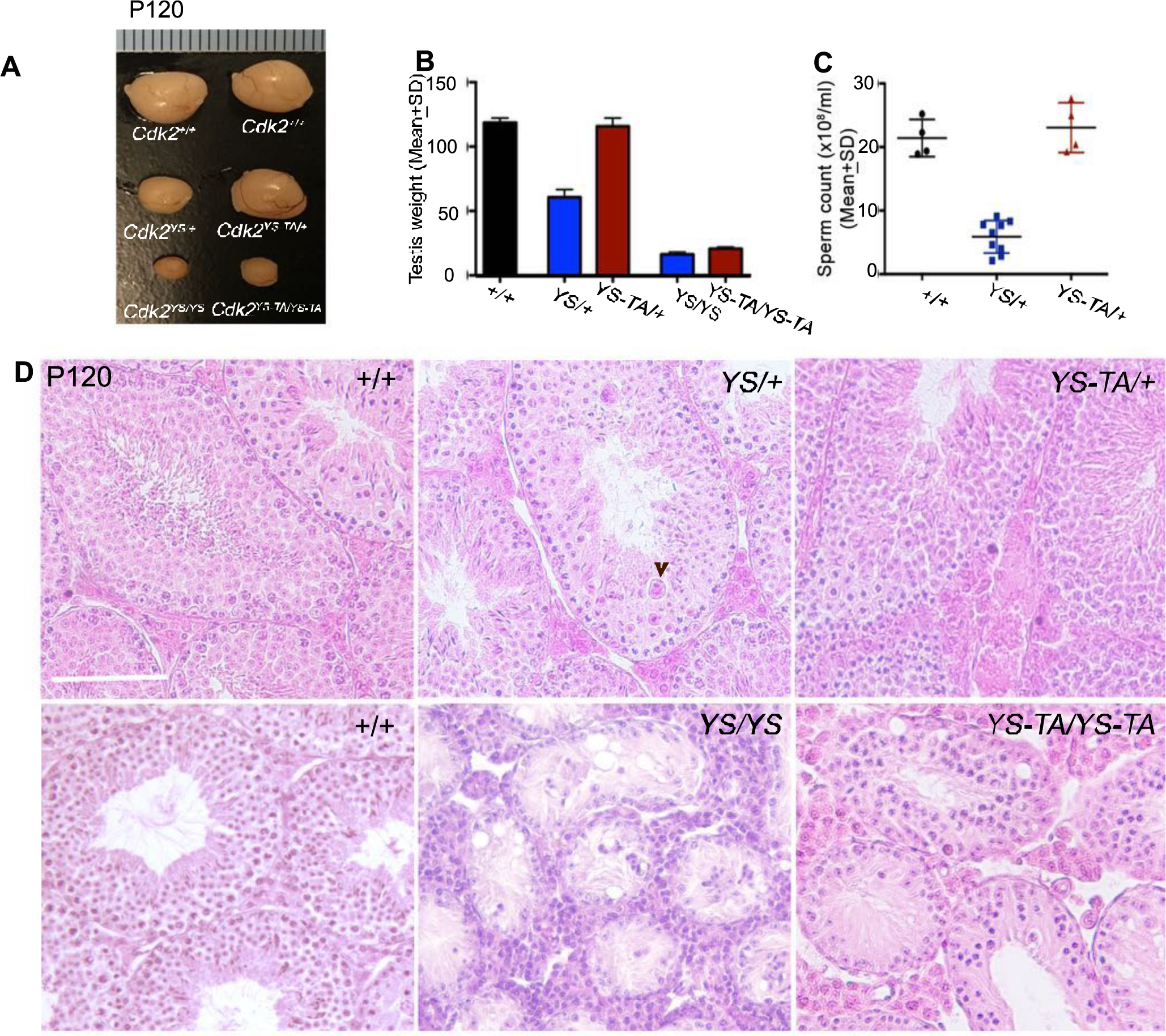
Phenotypic effects of CDK2 Tyr15 and Thr160 phosphorylation during spermatogenesis. **(A)** Images and **(B)** weights of P120 testes of indicated genotypes. **(C)** Sperm counts of age-matched genotypes at P120. **(D)** H&E-stained testis cross-sections of P120 animals revealing cellularity of seminiferous tubules. A degenerating cell, possibly multinucleate, is indicated by the arrowhead. Error bars in B and C represents ± SD. Scale bar in D, 100 µM. YS = *Cdk2^Y15S^*; TA = *Cdk2^T160A^*.

## DISCUSSION

The remarkable functional longevity of spermatogenesis in both mice and humans depends upon proper establishment and homeostasis of the SSC pool. A key event in male germline establishment is the seeding of prepubertal testes with a quiescent (G1-arrested) population of gonocytes (also called prospermatogonia). About 2-3 days after birth, these cells re-enter the cell cycle to expand and differentiate into SSCs that establish a permanent, renewable pool of cells that can initiate waves of spermatogenesis throughout life. Once established, there must be a fine balance of SSC self-renewal vs differentiation to maintain long term homeostasis and fertility.

SSC biology has been the subject of intense investigation. The timing and synchronization of the differentiation steps of spermatogenesis are well-described, and are controlled by exquisitely-timed production of retinoic acid (Endo et al., 2015). However, less is known about the molecular mechanisms underlying the gonocyte-to-spermatogonial transition (GST), the precise regulation of SSC renewal *vs* differentiation, and how infertility in people may result from disruption to these processes.

There is some debate as to the character of SSCs and their behavior with respect to cycling activity (Braun et al., 2018; Huckins, 1971a). In some tissues, the ability to continuously produce differentiated cells depends upon proper maintenance of a relatively infrequently-cycling population of stem cells, as in the case of the hematopoietic system. Overproliferation of hematopoietic stem cells (HSCs) caused by deregulated cell cycle control can lead to their exhaustion or transformation (Pietras et al., 2011). Similar to HSCs and most somatic stem cell niches, it has been hypothesized that there is a slow cycling population of SSCs that maintain the germline (Braun et al., 2018; Huckins, 1971b). However, there is also evidence for rapid turnover of SSCs (Klein et al., 2010) and for the ability of chains of differentiating spermatogonia (A_pr_ and A_al_) to fragment and de-differentiate to become A_s_ SSCs (Hara et al., 2014; Nakagawa et al., 2010). Regardless of various models proposed for the identity and behavior of “true” SSCs [reviewed in (de Rooij, 2017)], proper regulation of the cell cycle is essential. This is underscored by our studies, which implicate CDK activity regulation as being crucial for the GST and SSC renewal *vs* differentiation.

Phenotypes of certain mouse mutants have provided some insight into how regulation of cell cycle impacts spermatogonial maintenance and proliferation. Conditional germline knockout of the *Rb* tumor suppressor, a negative cell cycle regulator, causes infertility by abolishing the ability of SSCs to self-renew (Hu et al., 2013). In contrast to somatic stem cell systems in which *Rb* deficiency causes progenitor proliferation and a failure of terminal differentiation, the entire *Rb^-/-^* germ cell pool underwent a single round of spermatogenesis that yielded functional sperm (Hu et al., 2013). Cell cycle progression in normal cells requires inactivation of Rb by CDK/cyclin-mediated phosphorylation (including by CDK2/cyclinE), thus allowing expression of genes regulated by E2F transcription factors (Rubin, 2013). Moreover, ablation of *Plzf*, which negatively regulates the cell cycle by both inhibiting key regulators (McConnell et al., 2003; Yeyati et al., 1999) and also the self-renewal signal of GDNF (Hobbs et al., 2010), causes a less severe phenotype than Rb deficiency. *Plzf^-/-^* males are infertile due to a defect in SSC maintenance that leads to progressive germ cell loss and SCOS (Buaas et al., 2004; Costoya et al., 2004). Thus, a cell cycle-centric interpretation of these phenotypes is that unrestrained cycling (as in *Rb^-/-^*) causes efficient differentiation of all SSCs, but a moderate loss of cell cycle control (as in *Plzf^-/-^*) increases the propensity of SSCs to differentiate rather than self-renew. In the context of this model, CDK2^Y15S^ might have a lower impact on cell cycle control than *Rb* and *Plzf* mutants, possibly mediated by abnormal but partial inactivation of Rb. The result is abnormal SSC proliferation but only partial differentiation, ultimately leading to death of the aberrant cells (Clusters D and E) before meiotic entry. A mouse model in which both the Thr14 and Tyr15 negative phosphoregulatory sites were mutated to Ala and Phe, respectively (“*Cdk2^AF^* “), has previously been reported (Zhao et al., 2012). Although specific CDK2-associated kinase activity was not investigated in this mouse model, homozygous MEFs (mouse embryonic fibroblasts) exhibited accelerated entry into S phase. Interestingly, though no data was presented, the only significant defect in these mice was male infertility due to an apparent absence of germ cells in homozygotes and also in heterozygotes in one strain background (Zhao et al., 2012). In that regard, this model appears to resemble our *Cdk2^Y15S^* mouse. This might indicate that the Thr14 residue is redundant or not important for negative regulation of SSCs, although careful comparative phenotyping would need to be done.

Another explanation for the observed phenotypes of *Cdk2^Y15S^* cells might be informed by biochemical effects of similar mutations. Counterintuitively, while we observed elevated CDK2 kinase activity in spleens from heterozygous mice, the activity was greatly reduced in homozygous spleens. However, CDK2 kinase over-activation has been previously shown to be deleterious to cell cycle progression (Hughes et al., 2013). We postulate that, at least in spleen, cell autonomous regulation limits CDK2 activity within an acceptable range of activity to avoid this toxicity. Mutation of both negative phosphoregulatory sites in CDK2 (Thr14 and Tyr15) in human Hct116 cell lines caused premature S phase progression, DNA damage accumulation, and genomic instability leading to S phase arrest (Hughes et al., 2013). In these cells, degradation of Cyclin E was found to be increased and this could be relieved by p21 depletion, suggesting the presence of cellular feedback loops to attempt to reduce the levels of CDK2 activity upon premature activation (Hughes et al., 2013). In consideration of the aforementioned data of others, and our data showing elevated replicative activity and apoptosis of GFRa1^+^ cells in *Cdk2^Y15S/Y15S^* testes, we propose a model (Fig. 8) in which CDK2-associated kinase activity must be tightly regulated during specific stages of the cell cycle in SSCs, otherwise those cells attempting to differentiate will eventually die from cell cycle dysregulation.

**Figure 8:**
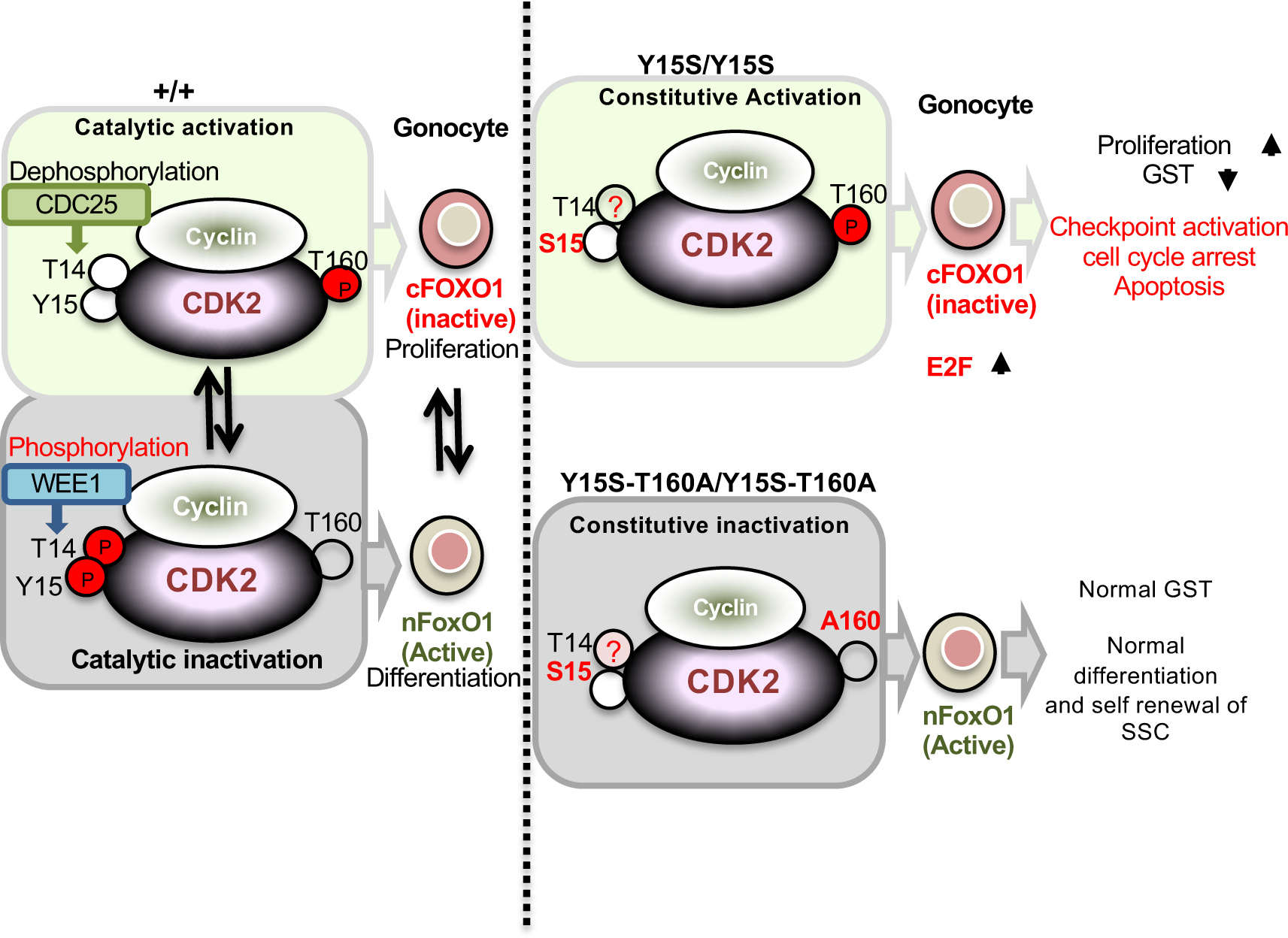
Model showing regulation of spermatogonia differentiation by Tyr15 and Thr160 residues of CDK2.

A caveat to the model is that it is based on cell cycle studies in cultured cells. Here, the major effect of the *Cdk2^Y15S^* mutation is on differentiation of germ cells residing in a complex milieu. Whereas there are mutations that reduce the number of perinatal gonocytes (typically stemming from PGC defects) (Hamer and de Rooij, 2018), the *Cdk2^Y15S^* phenotype is unique in that the GST is defective. Nevertheless, studies of other mutants give insight into how perturbations to cell cycle regulation impact gonocytes. As we described, *Cdk2^Y15S/Y15S^* germ cells fail to relocalize FOXO1 from the cytoplasm to the nucleus, a hallmark of the GST which normally occurs between P3 and P21 (Goertz et al., 2011; Kang et al., 2016; Pui and Saga, 2017). This cytoplasmic localization occurs as a result of CDK2-dependent phosphorylation (inactivation) of FOXO1 (Huang et al., 2006), which is essential for SSC maintenance and differentiation (Goertz et al., 2011). Since FOXO1 nuclear localization is disrupted in *Cdk2^Y15S^* mutants, we conclude proper cell cycle regulation and/or CDK2 kinase activity is a pre-requisite for the GST, and thus lies upstream in the developmental evolution/progression. Mice lacking the transcription factor *Glis3* partially resemble *Cdk2^Y15S^* mutants; they lack FOXO1 nuclear localization in neonatal gonad and fail to undergo a normal GST, as indicated by decreased expression of genes associated with the permanent pool of undifferentiated spermatogonia (Kang et al., 2016). However, it is unknown whether GLIS3 regulates these genes directly or indirectly, so its relationship to CDK2 in the developmental hierarchy is unclear.

Interestingly, the first round of spermatogenesis in *Cdk2^Y15S/Y15S^* mice occurs normally, although meiosis is still interrupted as in null mice. This first wave of spermatogenesis is claimed to arise directly from a subset of gonocytes that do not express *Ngn3* (Yoshida et al., 2006). We can conclude that differentiation of this subtype of gonocytes neither requires CDK2 nor is affected by its abnormal activity in *Cdk2^Y15S^* mutants. This underscores the caution needed when extrapolating data from cultured cells or from somatic cells (such as spleen) to germ cells, and even from one germ cell type to another.

These investigations into the *Cdk2^Y15S^* mouse stemmed from a project to investigate the genetic basis of human infertility, and involves modeling human SNPs in mice to determine possible pathogenicity (Singh and Schimenti, 2015). A lesson learned from this allele, and others that we have generated while pursuing this project (most not yet published) is that very few predicted deleterious variants are nulls, and that extensive phenotyping is required to understand their impact. Generally speaking, unless this mutation is entirely unusual in its effects, it raises the possibility that mice and people that present with non-obstructive azoospermia, and which have histopathology superficially resembling SCOS, may in fact have an occult pool of SSCs that have lost the ability to differentiate, but might be stimulated to do so after appropriate intervention. Additionally, as an example of an autosomal semi-dominant male infertility allele, it emphasizes the importance of considering monoallelic alterations when attempting to identify genetic causes of an individual patient’s infertility.

## AUTHOR CONTRIBUTIONS

P.S. conducted all experiments related to mouse breeding, cytology, histology, and preparing samples for sc sequencing. She also contributed to bioinformatics analyses and writing the paper. R.K.P. did most of the scRNA-seq analyses, with contributions from J.K.G. and supervision from A.G. N.P performed the kinase activity assays under the supervision of P.K., and they wrote the relevant sections and contributed to overall interpretations. D.A.P. made the initial observations that *Cdk2^YS/YS^* mutant adult testes were not devoid of spermatogonia. J.C.S. oversaw the entire study and shared the bulk of manuscript writing with P.S.

## ACKNOWLEDGEMENTS

This work was supported by a grant from the National Institutes of Health (R01 HD082568 to JCS and P50HD076210 to AG), and contract CO29155 from the NY State Stem Cell Program (NYSTEM). The authors would like to thank R. Munroe and C. Abratte of Cornell’s transgenic facility for generating the *Cdk2^T160A^* allele alone and in *cis* to the *Cdk2^Y15S^* change. Authors also thank the Proteomics Facility of Cornell University for providing the mass spectrometry data and NIH SIG 1S10 OD017992-01 grant support for the Orbitrap Fusion mass spectrometer. Confocal Imaging data was acquired in the Cornell BRC-Imaging Facility using the NIH-funded (S10OD018516) Zeiss LSM880 confocal/multiphoton microscope (u880).

## Materials and Methods

### Mouse strains and breeding

Mice used were on a mixed genetic background (FVB/NJ and B6(Cg)-Tyr^c-2J^/J). Experiments with the animals were performed under a protocol (2004-0038) approved by Cornell’s Animal Care and Use Committee. For female fertility tests, 8-10 wk-old WT or mutant homozygote females were mated with age-matched B6(Cg)-Tyr^c-2J^/J males.

### Production of CRISPR/Cas9-Edited Mice

The *Cdk2^Y15S-T160A^* allele was generated using CRISPR/Cas9 genome editing, essentially as described previously (Singh et al. 2015; Varshney et al. 2015) sgRNAs and ssODNs are listed in (Table S1). Briefly, *In vitro* transcription of sgRNA were performed using MEGAshortscript™ T7 Transcription Kit (Ambion; AM1354), and ssODN were availed from IDT. sgRNA, ssODN, Cas9 protein and mRNA (44758; Addgene) were co-microinjected into zygotes [F1 hybrids between strains FVB/NJ and B6(Cg)-Tyrc-2J/J] using the reagent concentrations listed in Table S1. Edited founders were identified either by subcloning followed by Sanger sequencing using primers and annealing temperatures listed in Table S1. For generating the *Cdk2^Y15S-T160A^* allele, *Cdk2^Y15S/Y15S^* females were used as embryo donors, thus, the Y15S alteration was not re-introduced by CRISPR.

### Testes histology and Immunohistochemistry

For histological analyses, testes were fixed for 24 h at room temperature (RT) in Bouin’s solution, paraffin-embedded, sectioned at 7 μm, and then stained with H&E. For immunohistology, testes were fixed in 4% paraformaldehyde for ∼24 h, paraffin-embedded, sectioned at 7 μm, and deparaffinized. Antigen retrieval for different antibodies was performed as indicated in Table S2. Sections were blocked in PBS containing 5% goat serum for 1 h at RT, followed by incubation with primary antibodies for 12 h at 4 °C and detection by the secondary antibody (see antibody details in Table S2).

### Whole-Mount Seminiferous Tubule Staining

Seminiferous tubule whole mounts were prepared as described previously (Savitt et al. 2012), with minor modifications. Briefly, seminiferous tubules were manually isolated in phosphate-buffered saline (PBS) and interstitial tissue was washed thoroughly followed by fixation in 2% paraformaldehyde for 2 h. After extensive washing, tubules were processed for detection of progenitor spermatogonial markers and EdU labeling. To detect ZBTB16, FOXO1, cPARP1 and LIN28, tubules were blocked in PBSS buffer (5% goat serum, 0.5 % Triton X-100 in 1x PBS) for 2 h at RT following primary antibody incubation for 12 h-18 h at 4°C in the same buffer. Tubules were washed at room temperature twice for 15 minutes, five times for 1 hour in PBSS, and then incubated overnight at 4°C with secondary antibodies. To detect GFRA1, the blocking and all incubations and washing steps are done exclusively in PBS-AT (1% BSA, 0.1% Triton X-100 in 1x PBS) buffer. Briefly, isolated tubules were blocked for 2 h and incubated in goat anti-GFRA1 antibody diluted in PBS-AT buffer. This was followed by five subsequent washing (30 min each) and detection by secondary antibodies. While performing co-immunolabeling of GFRA1 with other spermatogonial markers, ZBTB16, FOXO1, cPARP1 and LIN28 were always detected prior to GFRA1.

For EdU incorporation, mice were injected intraperitoneally with 100 mg/kg body weight of EdU (PY7562; Berry & Associates) before harvesting testes. Testes were then removed and immediately processed for whole-mount staining following the procedure described above, or fixed in 4% PFA for immunohistochemical staining. Following antibody staining, freshly prepared Click reaction cocktail (10mM (+)-sodium-L-ascorbate, 0.1mM 6-Caboxyfluorescein-TEG azide (FF 6110; Berry & Associates) or Alexa Fluor® 594 Azide (A10270; Invitrogen™), and 2mM copper (II) sulfate in water) was added into whole mount tubules or testes cross sections and removed quickly after 45 sec. Excess EdU was removed by extensive washings (2x 1h and 2x 12h at 4 °C) using PBS-AT buffer. Following washing, tubules were mounted in Vectashield (Vector Laboratories, Burlingame, CA). Click reaction was always performed following antibody detection. In each animal, we counted all A_s_, A_pr_ and A_al_ GFRa1^+^ cells positive or negative for EdU seminiferous tubules at least >8 mm in length.

### Imaging

Slide preparations were scanned and tiled images of testes cross sections or seminiferous tubule whole mounts were acquired using a Laser Scanning Confocal microscope (u880, Carl Zeiss, Germany) using Plan Apo 40× water immersion objective (1.1 NA) and Zen black software. Initial laser power adjustment were performed to avoid saturation of signal: argon laser-488 nm, blue-diode-405 nm, DPSS laser—561 nm and HeNe-633. Additional images of testes cross sections were obtained through an Olympus BX51 microscope with objectives 100×/1.35 NA infinity/0.17 or objective or 10×/0.3 NA, respectively using Olympus Cell Sens software (Olympus). Following identical background adjustments for all images, cropping, color, and contrast adjustments were made with Adobe Photoshop CC 2017.

### In vitro kinase assay

These were performed as described elsewhere (Lim et al. 2017). Briefly, 1ug of anti CDK2 antibody was added to 0.5µg of protein lysate and rotated overnight (approx. 16hrs) at 4°C. Twelve µl of protein A beads was added to each sample and left to mix for a further 2 hours. Prior to their use, these beads were washed twice previously using 1ml of EBN buffer (80 mM beta-glycerophosphate; 15 mM MgCl_2_; 20 mM EGTA-adjusted to pH7.3 with KOH; followed by adding 150mM NaCl, 0.5% NP40 to 0.and 1X protease inhibitor cocktail). After binding, the beads were spun down and washed with EBN buffer for 3 times. Pellets were resuspended in 14ul EBN buffer (containing protease inhibitors: cOmplete, Mini, 11836153001; Roche) followed by adding 6ul of Histone H1 (260ng/ul) and 9ul of kinase assay buffer (15mM EGTA, 25mM NaF, 250mM sodium beta glycerophosphate, 5mM DTT, 20mM MgCl_2_, 21uM ATP). Following 20 min incubation at room temperature, 1ul of ATP [γ-32P] containing a specific radioactivity of 5uCi (Perkin Elmer: #BLU502A) was added to each sample and left for 30 minutes at 30°C, 400rpm. 6X SDS sample buffer (2M beta-mercaptoethanol 0.375 M Tris pH 6.8.12% SDS,60% glycerol, 0.6M DTT,0.06% bromophenol blue) was added to a final concentration of 1X and each sample was boiled at 95°C for 5 minutes. 12ul of each sample were analyzed for Coomassie stained Histone H1 protein bands and Phosphosignal on a phosphor screen cassette for 6-24hrs. Phosphosignal was quantified using FLA7000 phosphimager (Fujifilm).

### Western blotting

Protein was extracted using EBN buffer (see above) with protease inhibitors. Samples were boiled at 95oC for 5 minutes and were separated by SDS/PAGE (12% acrylamide). Separated proteins were electrotransfer to 0.2uM nitrocellulose (Biorad #1620112) or PVDF membrane (Immobilon-P membrane, IPVH00010; EMD Millipore), and then blocked in 5% nonfat milk for 45 min at RT. Membrane was washed 3X 10 minutes in TBST (0.14M NaCl, 15mM KCl, 25mM Tris Base, 1% Tween20) at room temperature followed by ∼16 h incubation with primary antibody, washing, and ∼1 h incubation with HRP conjugated secondary antibody (as stated in Table S4) for 45 minutes at room temperature. Signal was detected using Luminata Classico Western HRP substrate (WBLUC0100; EMD Millipore). Densiometric analysis of western blot bands was performed using Fujifilm Multi Gauge software Ver. 3.1.

### Site directed mutagenesis

Site-directed mutagenesis for obtaining CDK2-Y15F and CDK2-Y15S cDNA was accomplished using the QuikChange Lightning Site-Directed Mutagenesis Kit (Agilent, La Jolla, CA; Cat# 210518) using primers sets mentioned in Table S3. All mutations were confirmed by Sanger sequencing.

### CDK2 overexpression and immunoprecipitation

The (Myc-DDK-tagged)-CDK2 cDNA was obtained from Origene (Cat#RC200494). Y15F and Y15F mutant cDNA were generated as described above and transfected into HEK-293T cells (ATCC; CRL-3216) using TransIT®-LT1 Transfection Reagent (Mirus; MIR2305) following user instructions. Lysates were prepared from harvested cells in lysis buffer (50mM Tris pH7.5, 150mM NaCl, 0.5% Triton X-100, 5 mM EDTA) containing phosphatase (Thermo Scientific; 78428) and protease inhibitors (Roche; 04693159001). 400 µg of each lysate was immunoprecipitated using c-Myc-Tag IP/Co-IP Kit (Pierce, 23620). Proteins were separated on 12% poly acrylamide gel by SDS-PAGE and CDK2 protein was visualized by western blot analysis.

### Protein Identification by nano LC/MS/MS Analysis

In-gel trypsin digestion of immunoprecipitated CDK2 protein was performed as described earlier (Yang et al. 2007). The tryptic digests were subjected to nanoLC-ESI-MS/MS analysis on Orbitrap Fusion^TM^ Tribrid^TM^ (Thermo-Fisher Scientific, San Jose, CA) mass spectrometer equipped with a nanospray Flex Ion Source, and a Dionex UltiMate3000RSLCnano system (Thermo, Sunnyvale, CA) following a protocol described earlier (Yang et al. 2018; Thomas et al. 2017). The column was re-equilibrated with 0.1% FA for 23 min prior to the next run. For data-dependent acquisition (DDA) analysis, the instrument was operated using FT mass analyzer in MS scan to select precursor ions followed by 3 second “Top Speed” data-dependent CID ion trap MS/MS scans at 1.6 m/z quadrupole isolation for precursor peptides with multiple charged ions above a threshold ion count of 10,000 and normalized collision energy of 30%. MS survey scans at a resolving power of 120,000 (fwhm at m/z 200), for the mass range of m/z 375-1575. Dynamic exclusion parameters were set at 30 s of exclusion duration with ±10 ppm exclusion mass width. All data were acquired under Xcalibur 3.0 operation software (Thermo-Fisher Scientific).

### Mass spec data analysis

The DDA raw files for CID MS/MS were subjected to database searches using Proteome Discoverer (PD) 2.2 software (Thermo Fisher Scientific, Bremen, Germany) with the Sequest HT algorithm. All 3 raw MS files for three samples were used for database search. Processing workflow for precursor-based quantification. The PD 2.2 processing workflow containing an additional node of Minora Feature Detector for precursor ion-based quantification was used for protein and ptms identification. The database search was conducted against a *mouse* database containing ∼20153 entries downloaded from NCBI on Jan. 12, 2018 plus some common contaminants (246 entries) database. Two-missed trypsin cleavage sites were allowed. The peptide precursor tolerance was set to 10 ppm and fragment ion tolerance was set to 0.6 Da. Variable modification of methionine oxidation, deamidation of asparagines/glutamine, phosphorylation of serine, threonine and tyrosine and fixed modification of cysteine carbamidomethylation, were set for the database search. Only high confidence peptides defined by Sequest HT with a 1% FDR by Percolator were considered for the peptide identification. The final protein IDs contained protein groups that were filtered with at least 2 peptides per protein.

### Sperm counting

Sperm counting was done as described earler (Singh and Schimenti 2015).

### scRNA seq sample preparation

3.5 DPP testes were decapsulated in HBSS (Mediatech Inc.) and digested with 0.642 ml of Trypsin/EDTA (0.25%, Invitrogen Inc.) and 0.071 ml DNase I (1 mg/ml in HBSS, Sigma Inc.) at 37 °C for 3-5 min min. Digestion was stopped by adding 0.5 ml of media (HBSS + 10% FBS), the cells were filtered through a 70-µm cell strainer and resuspended in HBSS with 0.04% FBS. Single-cell 3’ RNA-seq sequencing libraries for Illumina were constructed using a 10X Genomics Chromium instrument, using the Chromium Single Cell 3’ Reagent kits (v2) following the manufacturer’s protocols. The final libraries were quantified by digital PCR and sequenced on Illumina sequencers, using an Illumina NextSeq500/550 75 bp kit (26 bp + 8 bp index read + 58 bp). The target number of cells captured from the input, single-cell suspension was 8700. The data were demultiplexed and aligned to the reference genome using the 10X Genomics Cellranger software (v2.2) and visualized using the 10X Genomics Loupe Cell Browser software (v2.0) packages.

### scRNA-seq data analysis

The raw scRNA-seq data was analyzed using cell ranger from 10X platform to generate a matrix of raw read counts, which was further analyzed in R using Seurat (Satija et al. 2015; Butler et al. 2018) and Monocle (Qiu et al. 2017). A cluster of germ cells from WT and YS were separated from rest of the testis cells based on *Ddx4* gene expression. We then isolated cell barcodes that had more than 10,000 UMIs per cells (see Results). Raw read counts for these selected germ cells were further filtered to keep only those genes that are detected in at least 10% of the cells. This resulted in an expression matrix of <u>10451 genes across 141 cells</u> (WT=69; YS=72). Due to high rate of drop-outs in mRNA-capture in scRNA-seq approaches, the expression values for mid- and lowly-expressed genes are often unreliable due to missing information. Hence, we recovered expression for each gene using SAVER (Huang et al. 2018), an approach that estimates gene expression from UMI-based scRNAseq data using information across all genes and cells. This estimated gene expression was used for further analysis. The most variable genes across all cells were identified using principle component analysis (PCA). Specifically, PCA was performed using log-transformed counts-per-million (CPM) expression values. The top 300 genes with highest absolute loading for PC1, top 300 genes for PC2 and top 300 genes for PC3 were chosen, which resulted into 744 unique genes. Furthermore, this list of variable genes was filtered to remove mitochondrial genes (n=10). The 141 germ cells were clustered into five groups using “Ward.D2” method based on expression estimates for the 734 highly variable genes. The pseudotime and trajectory analysis were performed based on the expression of 734 most variable genes using Monocle R code.

### Statistics

P values were calculated from unpaired Student’s t-test for all IHC and whole mount experiments.

