## Supplemental Figures for "CDK2 kinase activity is a regulator of male germ cell fate"

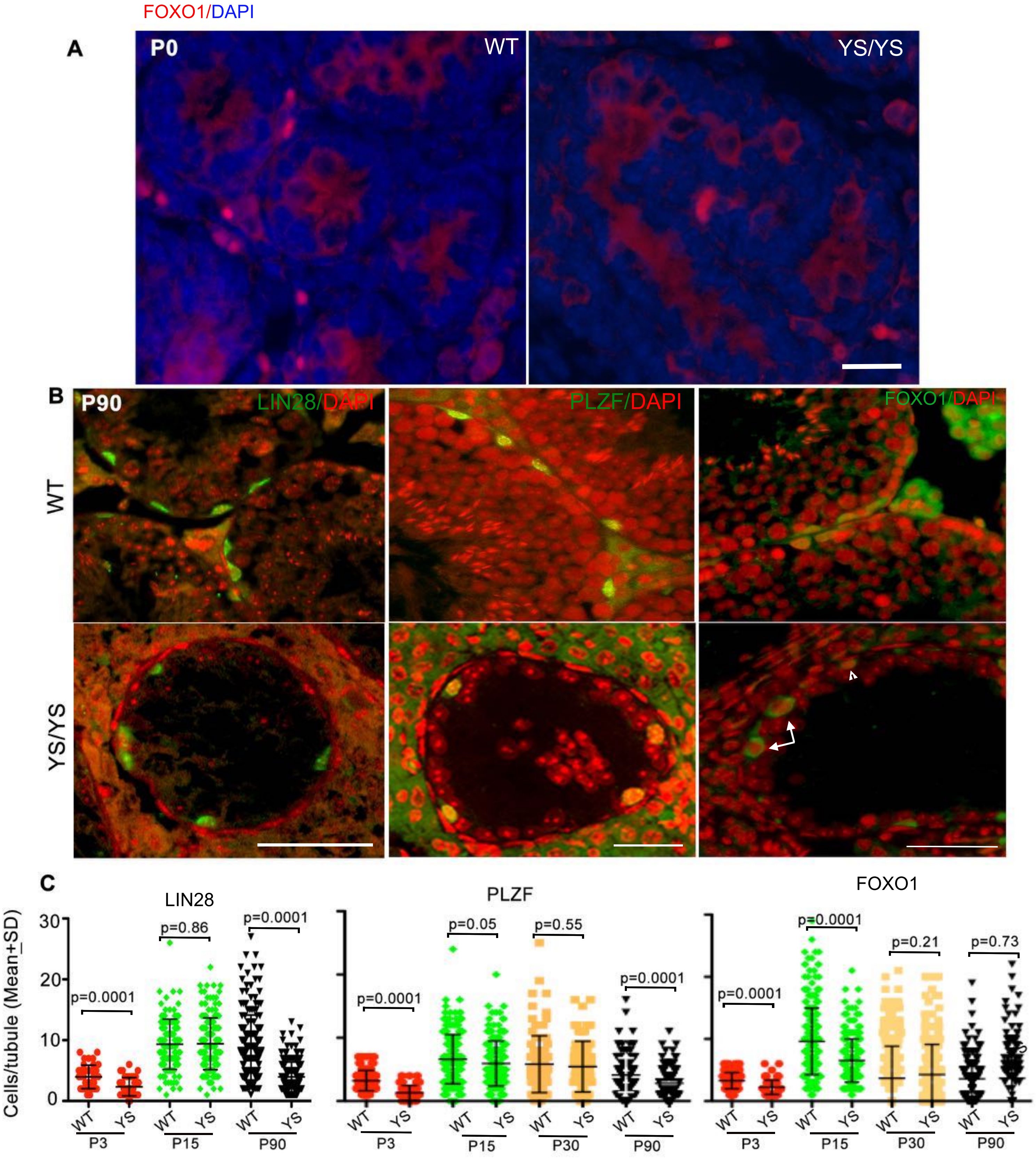


**Figure S1**: **Loss of phospho-CDK2 (Y15) can maintain undifferentiated spermatogonia that areunable to differentiate**. **(A)** PO testes cross sections stained with FOXO1 (red). **(B)** Testes crosssections of WT and mutant P90 gonads immunolabeled with indicated antibodies (all green). **(C)** Quantification of data in B at indicated ages and genotypes. FOXO1 quantification represent cyplasmic and nuclear FOXO1$^{+}$ spermatogonia. Arrow, cFOXO1; Arrowhead, nFOXO1. Nuclei are counterstained with DAPI (pseudo-colored red). Scale Bar, 20 μm in A and 50 μm in B.

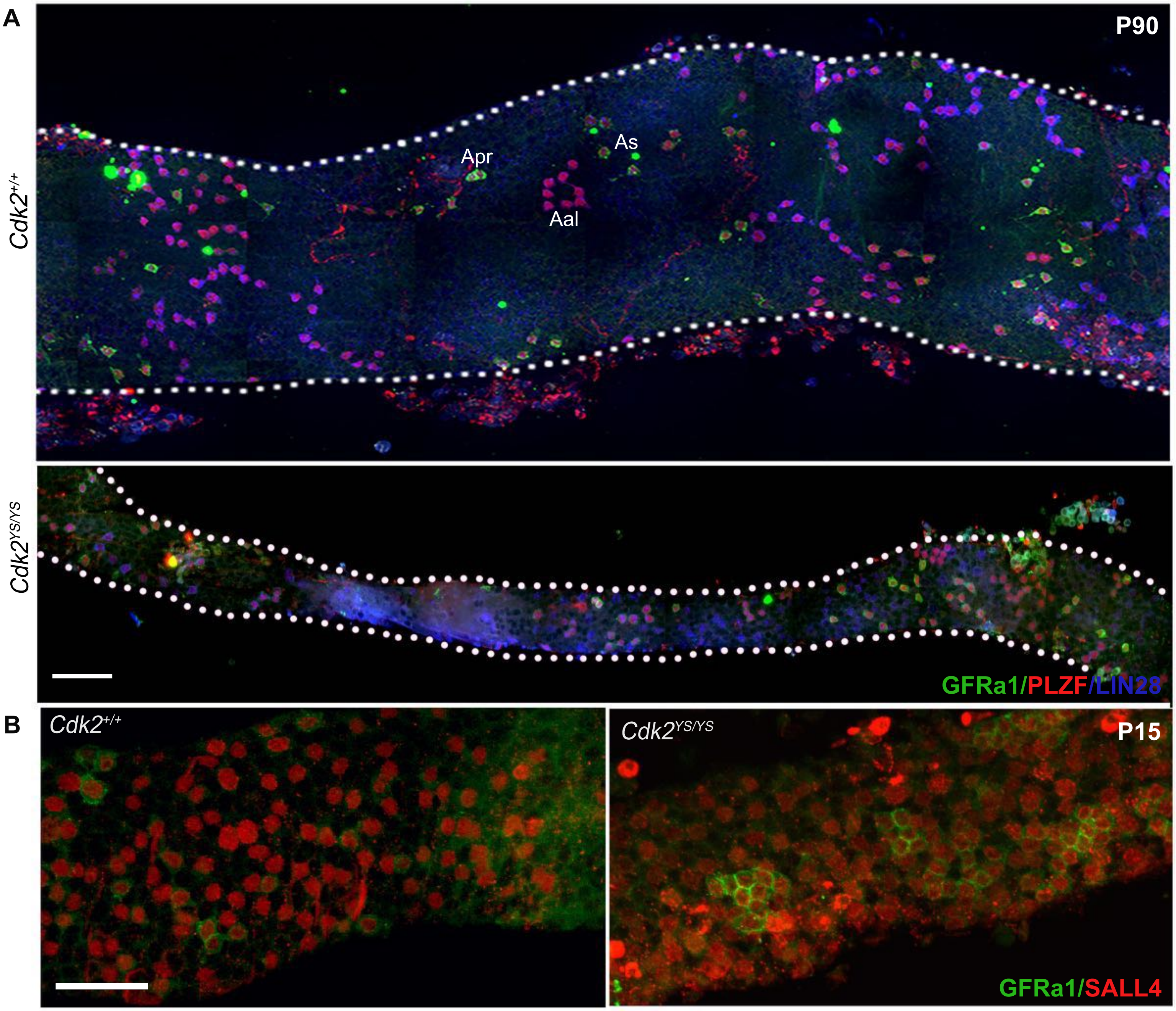


**Figure S2: (A)** Whole mounts of P90 seminiferous tubules stained with all markers of undifferentiated spermatogonia **(B)** Example of GFRa1 and SALL4 expressing clones in P15 tubules. Scale Bar, 50μm.

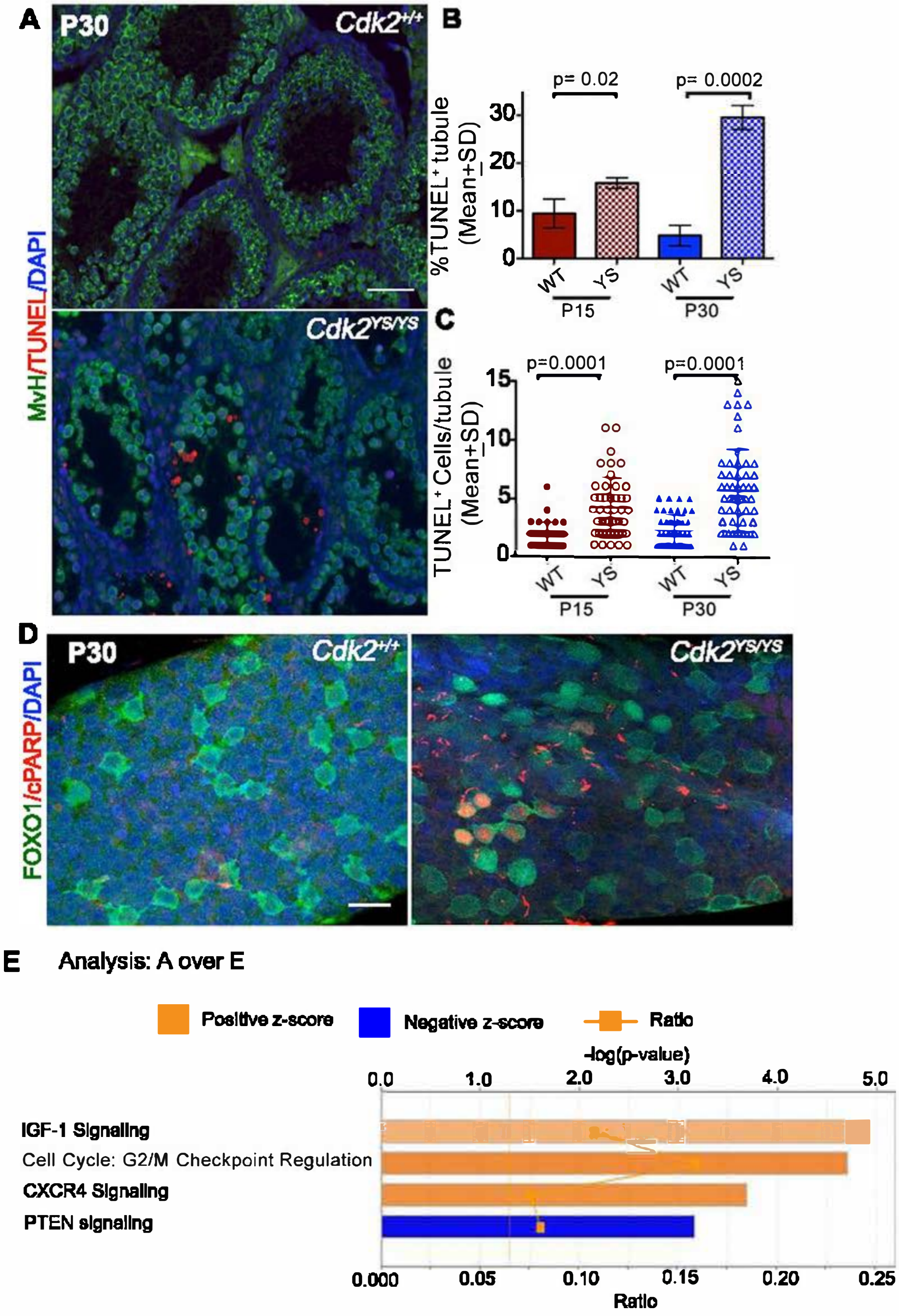


**Figure S3: Progressive loss of germ cells in *Cdk2*$^{YS/YS}$ mutant gonads**. **(A)** testis cross-section showing presence of TUNEL+ cells in MVH labeled (green) seminiferous tubule lumen of *Cdk2*$^{YS}$ homozygote. Bar diagrams representing percentages of TUNEL$^{+}$ tubules **(B)** and TUNEL$^{+}$ cells/tubule **(C)** at indicated ages and genotypes. **(D)** Double immunolabeling of WT and mutant seminiferous tubule with cleaved PARP (cPARP, red) and FOXO1 (green). **(E)** Ingenuity pathway analysis (IPA) on differentially expressed genes cluster A vs E showing predicted activation of indicated pathways in YS gonads. Scale Bar, 20 μm in D and 50 μm in A.

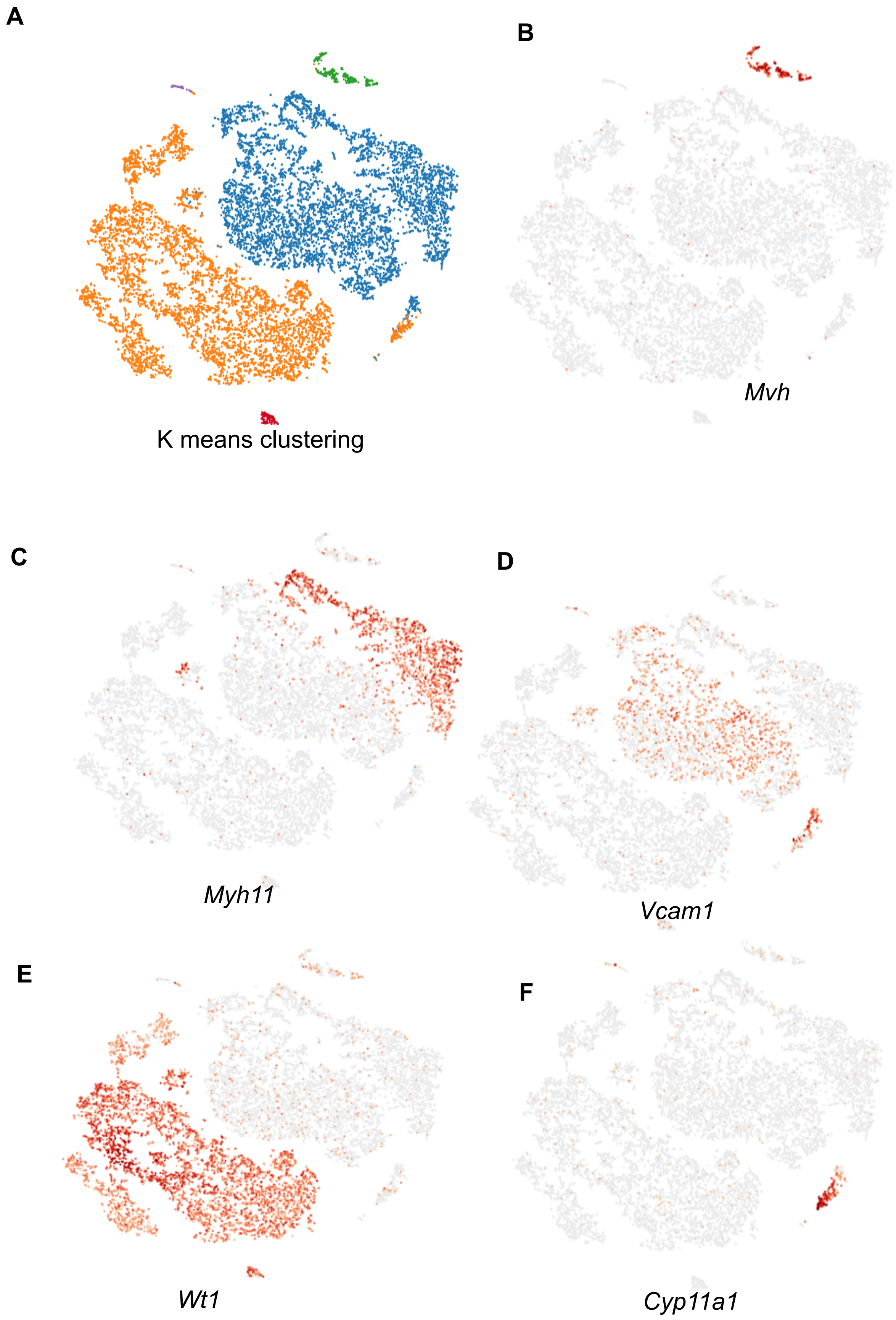


**Figure S4. The t-distribute stochastic neighbor embedding (t-SNE) plot identifies 5 clusters of spermatogenic based on K means (A).** Cells at different developmental stages are shown in different colors.**(B-F)** Corresponding developmental stages confirmed by specific expression signatures.

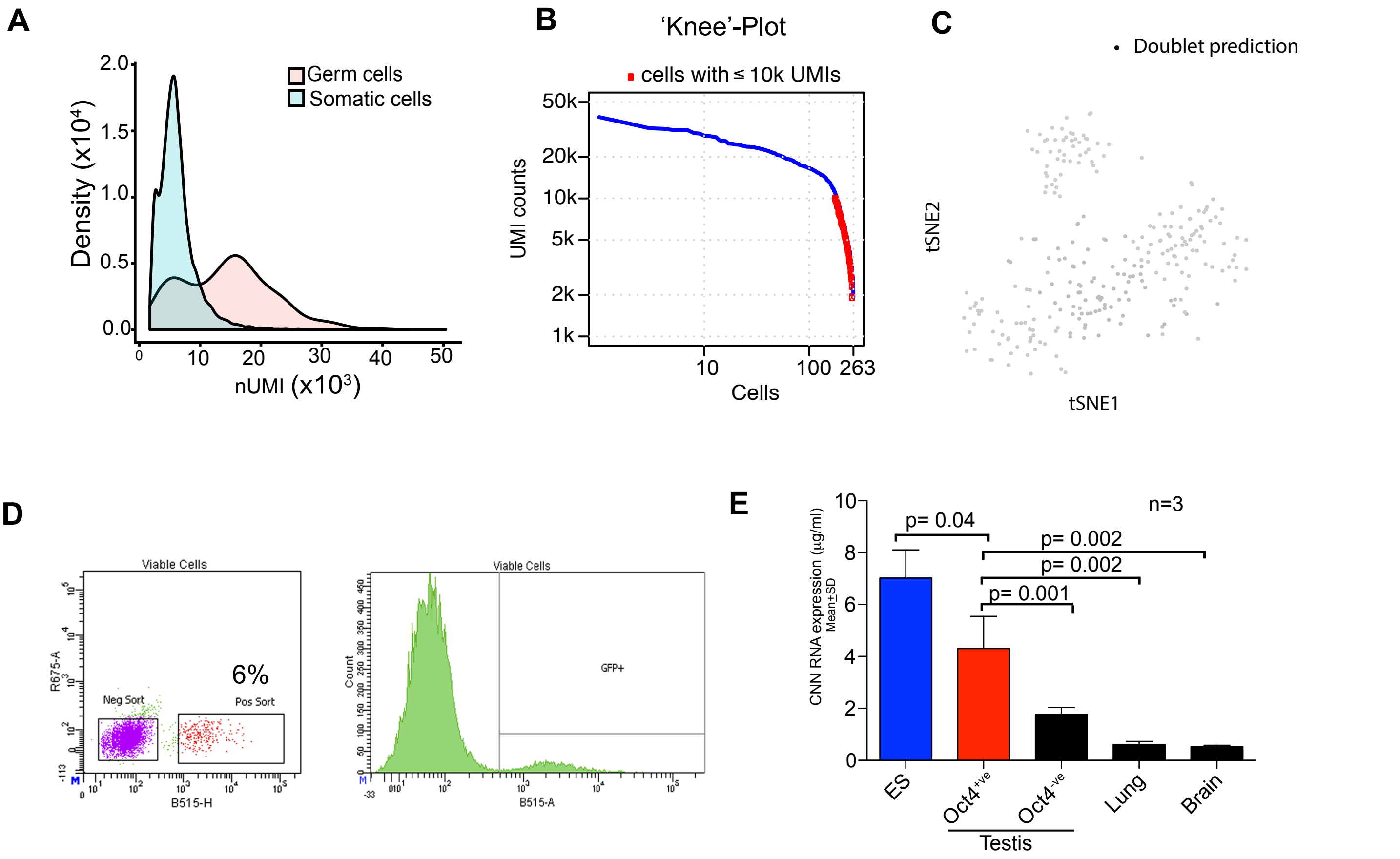


**Figure S5. Neonatal germ cells express more transcripts than somatic cells. (A)** nUMI distribution in germ and somatic cells. **(B)** The 'knee'-plot analysis showing that the cells with lower nUMIs are under the "knee", which arelikely to be artifacts **(C)** tSNE plot showing outcomes from Doublet- finder of germ cells of all three genotypes. Each dot represents analysed germ cells. Outcomes as black dot indicates doublet (0 doublets in 263 cells 0.0% cross-type doublet rate) **(D)** FACS analysis of Oct4-gfp+ and Oct4-gfp cells from neonatal WT gonad. **(E)** Quantification of total RNA per cell in P3-P5 testicular germ and somatic cells. CNN, Cell number normalized.

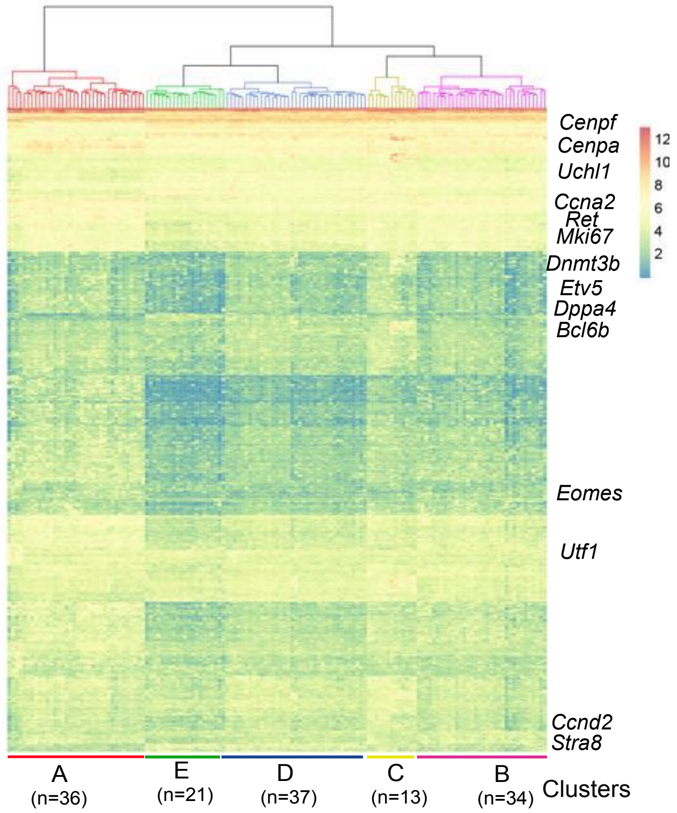


**Figure S6:** Hierarchical clustering of 141 germ cells from *Cdk2*$^{+/+}$ (69 cells) and *Cdk2*$^{Y15S/Y15S}$ *(*78 cells) gonads based on the expression (heatmap) of most divergently expressed genes resulted in five distinct clusters of cells. Bottom panel shows fraction of cells of indicated genotype in
each cluster. The cells from *Cdk2*$^{+/+}$ are shown as green, whereas those from *Cdk2*$^{Y15S/Y15S}$ are in red. Notable marker genes are highlighted to the right of the heatmap. Color key from green to red represents the relative (normalized and gene-wise Z scored) gene expression levels from low to high, respectively.

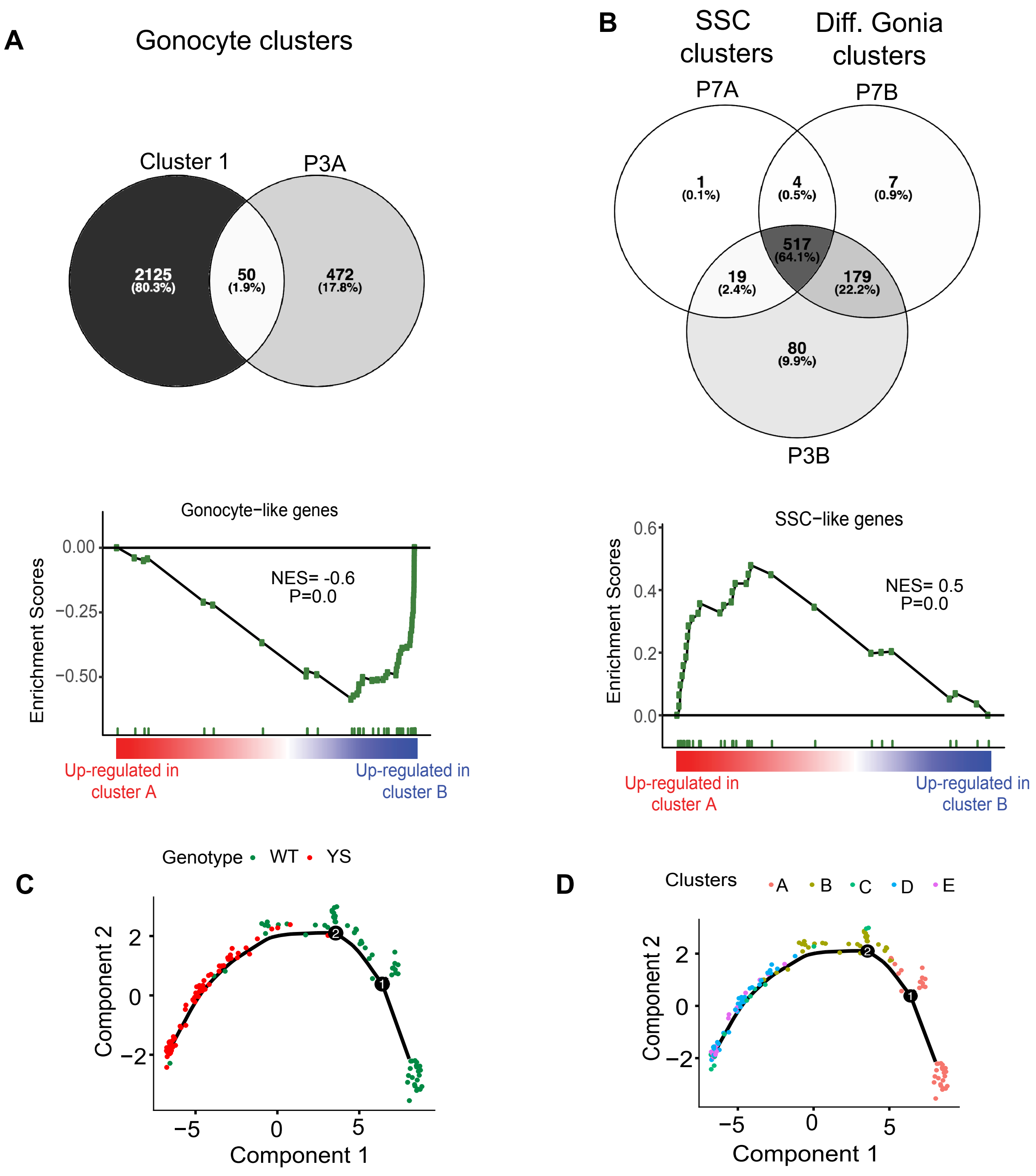


**Figure S7: Identification of germ cell subsets in *Cdk2*$^{+/+}$ and *Cdk2*$^{YS/YS}$ testes (P3)** Gene set enrichment analysis on **(A)** 50 overlapping genes defining gonocyte clusters and **(B)** 100 unique genes defining SSC clusters from publised datasets (Liao et. al., 2018; Song et al., 2016). **(C-D)** Monocle pseudotime trajectory analysis on data without SAVER-based imputations outlining the developmental timeline of germ cell subset clusters from WT and YS gonads defined in A,B and figure 5.

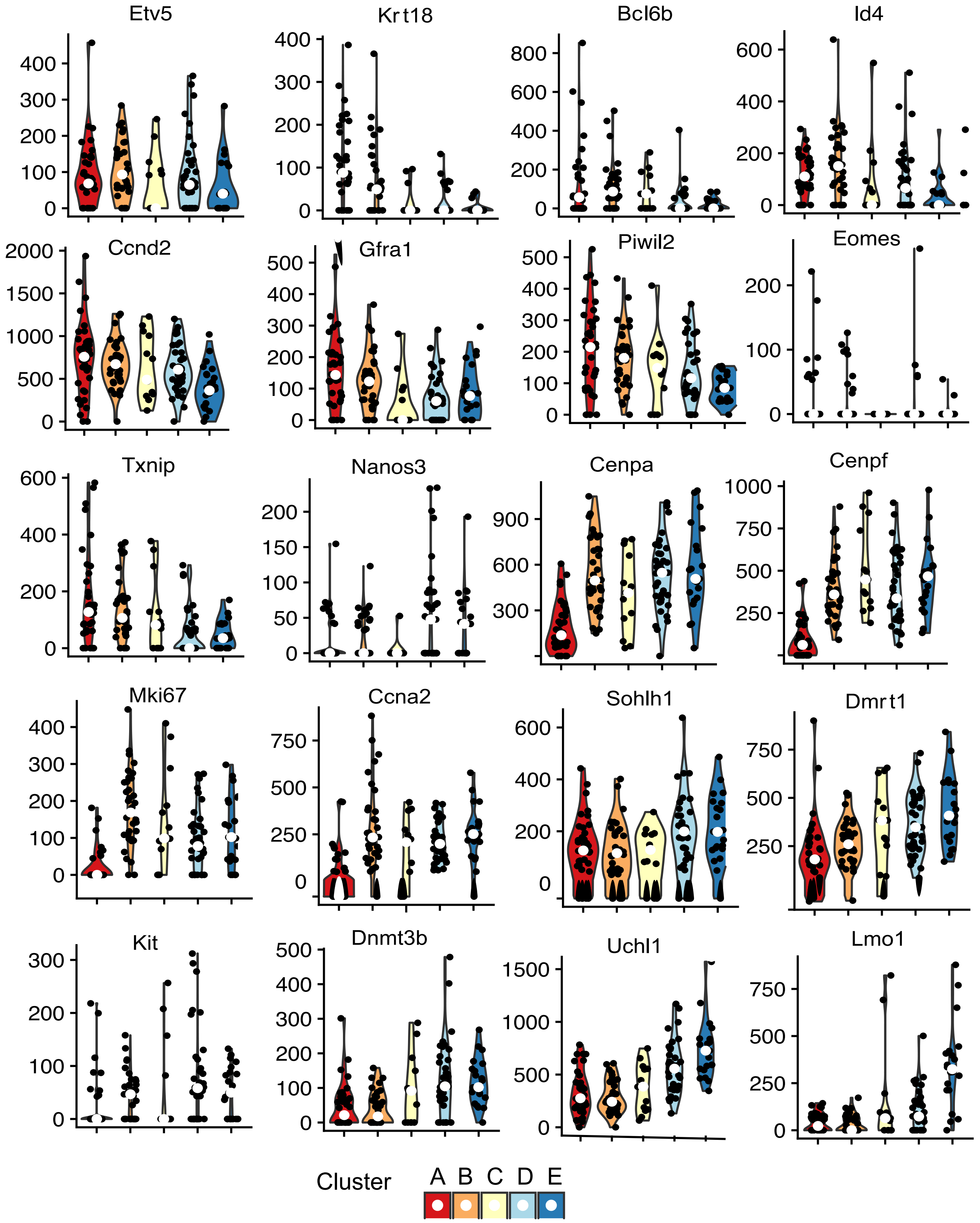


**Figure S8:** Violin plots of raw data (without SAVER- based imputation) showing expression of selected genes in different cell clusters.

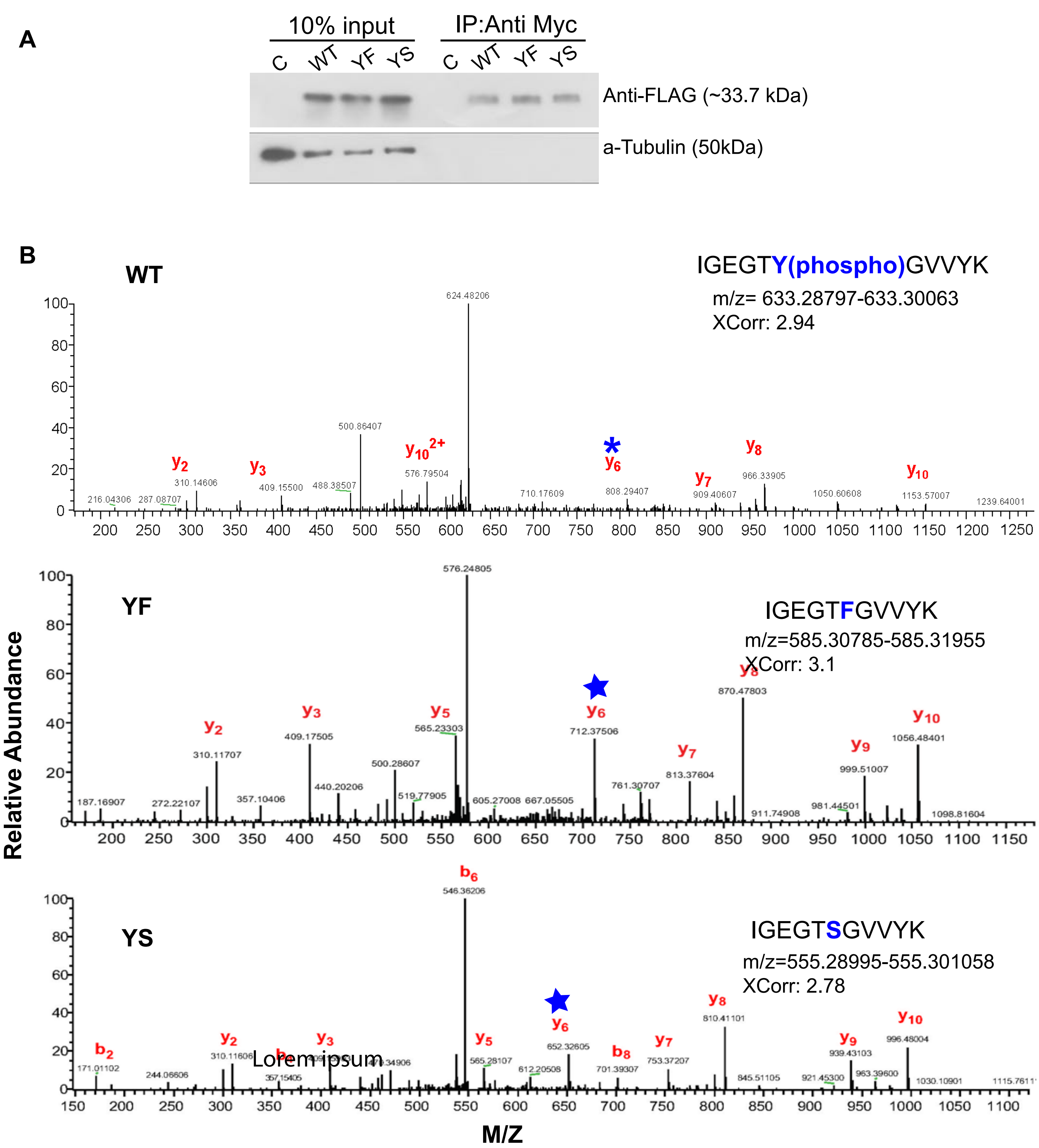


**Figure S9: Identification of phosphorylation status of Y15 and S15 residues of CDK2.** **(A)** Western blot showing overexpression and immunoprecipitation of WT and mutant CDK2. FLAG Myc-CDK2 cDNA was transfected into HEK293T cells, and immunoprecipitated and probed with anti-Myc and anti-FLAG antibodies, respectively. **(B)** MS/MS spectra showing identification of phosphorylation at Y15 residue only (WT condition), while phosphorylation is not detected in F15 or S15 transgenes. Asterisk (*) indicates ions that support the phospho-rylation of Y15 residue in WT. Star (★) represents ions indicating absence of phosphorylation in F15 and S15 residues.

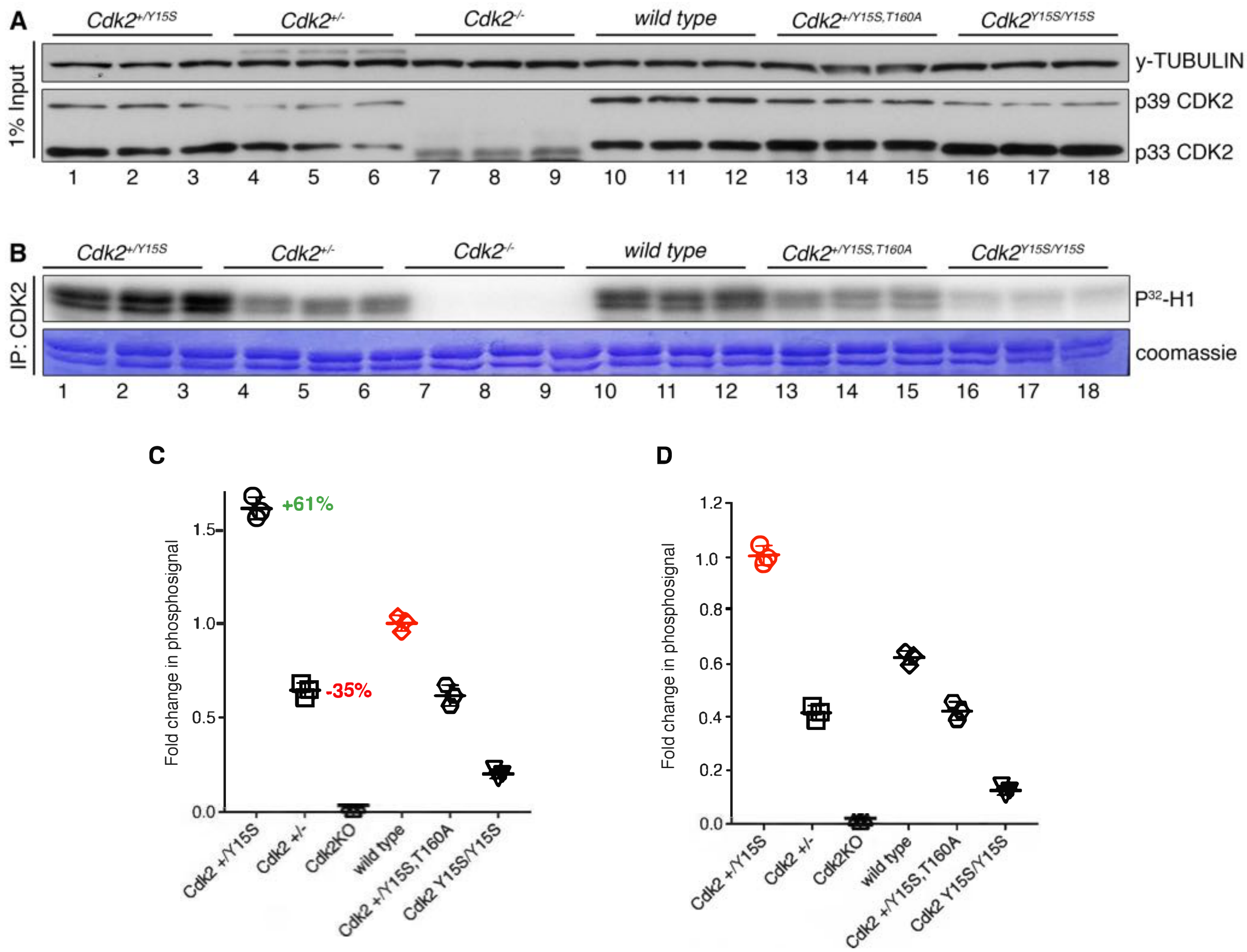


**Unpaired T-tests**
**[Foldchange normalised to *wild type*]** *wild type* VS *Cdk2*$^{+/Y15S}$ P=0.0001
*wild type* VS *Cdk2*$^{+/-}$ P=0.0004
*wild type* VS *Cdk2*$^{-/}$ P<0.0001
*wild type* VS *Cdk2*$^{+/Y15S,T160A}$ P=0.0013
*wild type* VS *Cdk2*$^{Y15S/Y15S}$ P<0.0001

**Unpaired T-tests**
**[Foldchange normalised to *Cdk2*$^{+/Y15S}$]**
*Cdk2*$^{+/Y15S}$ VS *wild type* P=0.0001
*Cdk2*$^{+/Y15S}$ VS *Cdk2*$^{+/-}$ P<0.0001
*Cdk2*$^{+/Y15S}$ VS *Cdk2*$^{-/-}$ P<0.0001
*Cdk2*$^{+/Y1SS}$ VS *Cdk2*$^{+/Y15S-T160A}$ P<0.0001
*Cdk2*$^{+/Y15S}$ VS *Cdk2*$^{Y15S/Y15S}$ P<0.0001

**Figure S10. Quantification of CDK2 kinase activity in spleens of *Cdk2* mutant mice. (A)** Western blots of 1% (10ug) of total protein lysate used for immunoprecipitation (input). Immunoblotting was performed using the specified antibodies on whole spleen lysates of *Cdk2*$^{+/Y15S}$ (1-3),*Cdk2*$^{+/-}$ (4-6),*Cdk2*$^{-/-}$ (7-9), wild type (10-12), *Cdk2*$^{+/Y15S-T160A}$ (13-15) or *Cdk2*$^{Y15S/Y15S}$ (16-18) mice. All spleens were extracted from 10 days old mice . **(B)** Kinase assays of CDK2-immunoprecipitates from spleen lysates shown in panel A. CDK2 immunoprecipitation was performed against 1 mg of whole spleen lysate using 1ug of anti-CDK2 antibody. The fold change in phosphosignal for each sample was calculated relative to the mean wildtype **(C)** or mean *Cdk2*$^{+/Y15S}$ **(D)** as displayed. Statistical tests were performed via unpaired-t test between each genotype.
