## Supplemental Table 3 for "CDK2 kinase activity is a regulator of male germ cell fate"

**Table S3. Oligonucleotides used for CRIPSR editing and genotyping**

|  | Gene name | | Sequence (5’-3’) |
| --- | --- | --- | --- |
| gRNA oligo | *Cdk2^T160A^* | | GAAATTAATACGACTCACTATAGG**TCATGAGTGTAAGTTCGGAC**GTTTTAGAGCTAGAAATAGC |
| Reverse primer | For gRNA template | | GCACCGACTCGGTGCCACTTTTTCAAGTTGATAACGGACTAGCCTTATTTTAACTTGCTATTTCTAGCTCTAAAAC |
| HR ssODN | *Cdk2^T160A^* | | CTGGCAGACTTTGGACTAGCAAGAGCCTTTGGAGTC CCA GTC CGTACTTATGCTCATGAGGTAAGTCCCTTTATGGTTTTCTTTTACAGT |
| Genotyping primers | *Cdk2^Y15S^*  (55^o^C) | F | CTTAAGCGCCACAACTTC |
|  |  | R | TCTCCTTGCGTTCCATC |
|  | *Cdk2^T160A^*  (60^o^C) | F | AAGACCCTTGGGTTGTCAGA |
|  |  | R | GAAACAAACCTCCCCCTCTC |
|  | Oct4-GFP  (56^o^C) | F | AAGTTCATCTGCACCACCG |
|  |  | R | TCCTTGAAGAAGATGGTGCG |
| Site directed mutagenesis primers | SDMYF-F  (52^o^C) | F | CAAAAGGTGGAGAAGATTGGAGAGGGAACATTCGGAGTGGTGTACAAAGCCAAAAAC |
|  | SDMYS-F  (52^o^C) | F | CAAAAGGTGGAGAAGATTGGAGAGGGAACATCCGGAGTGGTGTACAAAGCCAAAAAC |
|  | SDMYF-R  (52^o^C) | R | GTTTTTGGCTTTGTACACCACTCCGAATGTTCCCTCTCCAATCTTCTCCACCTTTTG |
|  | SDMYS-R  (52^o^C) | R | GTTTTTGGCTTTGTACACCACTCCGGATGTTCCCTCTCCAATCTTCTCCACCTTTTG |
