## Supplemental Table 4 for "CDK2 kinase activity is a regulator of male germ cell fate"

**Table S4. List of antibodies used in this study**

| Antibody name (Host) | Application | Cat# | Antigen retrieval method | Concentration |
| --- | --- | --- | --- | --- |
| FOXO1 (Rb) | IHC, WM | Cell signaling technology; 2880 | TE-pH9 | 1:100 |
| CDK2 (Ms) | Western | SantCruz; sc6248 | NA | 1:400 |
| CDK2 (Rb) | IP | PRK lab | NA | 1μg/IP |
| PLZF (Ms) | IHC, WM | Active Motif; 05313002 | TE-pH9 | 1:100 |
| SALL4 (Ms) | WM | Abcam; ab57577 | Na citrate-pH6 | 1:100 |
| LIN28 (Rb) | IHC, WM | Abcam; ab46020 | TE-pH9 | 1:200 |
| MVH (Ms) | IHC | Abcam; ab27591 | Na citrate-pH6 | 1:400 |
| GFRA1 | WM | R&D systems; AF560 | NA | 1:500 |
| Cleaved PARP | WM | BD Horizon^TM^; 564129 | NA | 1:100 |
| DYKDDDK Tag (Flag Tag) (Rb) | Western | Cell signaling technology; 14793 | NA | 1:1000 |
| α-Tubulin | Western | Abcam; ab11320 | NA | 1:800 |
| α-Tubulin | Western | Sigma Aldrich: T9026 | NA | 1:1000 |
| Goat anti-Mouse-488 | IHC, WM | Invitrogen; A11001 | NA | 1:1000 |
| Goat anti-Rabbit-594 | IHC, WM | A11012 | NA | 1:1000 |
| Donkey anti-Goat-488 | IHC, WM | A11055 | NA | 1:1000 |
| Donkey anti Rabbit-405 | WM | Jackson Immunoresearch; 711-025-152 | NA | 1:300 |
| Donkey anti Rabbit-594 | WM | Jackson Immunoresearch; 711-586-152 | NA | 1:400 |
| Donkey anti Mouse-594 | WM | Jackson Immunoresearch; 715-586-150 | NA | 1:400 |
| Goat anti-rabbit-HRP | Western | Pierce; 31462 | NA | 1/10,000 |
| Goat anti-mouse-HRP | Western | Pierce; 31432 | NA |  |

WM= Whole mount; IHC= Immunohistochemistry; IP= Immunoprecipitation
